# The discovery of 5mC-selective deaminases and their application to ultra-sensitive direct sequencing of methylated sites at base resolution

**DOI:** 10.1101/2024.12.05.627091

**Authors:** Weiwei Yang, Yan-Jiun Lee, Rebekah M. B. Silva, Amanda DeLiberto, Colleen Yancey, Daria McCallum, Jackson Buss, Rey Moncion, Jennifer Ong, Megumu Mabuchi, Dave Hough, Peter R. Weigele, Laurence M. Ettwiller

**Affiliations:** New England Biolabs Inc., 240 County Road, Ipswich, MA 01938, United States

**Author notes:** These authors contributed equally to this work (Weiwei Yang, Yan-Jiun Lee).

## Abstract

Mining phages for new enzymatic activities continues to be important for the development of new tools for biotechnology. In this study, we used MetaGPA—a method linking genotype to phenotype in metagenomic data—to identify deoxycytidine deaminases, a protein family highly associated with cytosine modifications in metaviromes. Unexpectedly, a subset of these deaminases exhibited a preference for 5-methylcytosine (5mC) over cytosine (C) in both mononucleotide and single-stranded DNA substrates. In a methylome sequencing workflow, preferential deamination of 5mC by these enzymes enabled direct conversion of methylated cytosine while completely eliminating any background deamination of unmodified cytosine. This direct conversion allows for precise identification of methylated sites at single-base resolution with unmatched sensitivity enabling broad applications for the simultaneous sequencing of genome and methylome.

## Introduction

Uncovering the molecular mechanisms behind the arms race between bacteria and their viruses has provided insights into deep evolutionary origins of eukaryotic innate immunity and epigenetics (Burroughs et al. 2015, Iyer et al. 2009). Components of the bacterial-phage defense and counter defense systems such as Type II restriction endonucleases, T4 polynucleotide kinase, and CRISPR-Cas enzymes have become indispensable tools for biotechnology (van der Oost and Patinios 2023). The discovery of such enzymes has recently been accelerated by the integration of high-throughput sequencing, comparative genomics, low-cost DNA synthesis, and functional screening. This multidisciplinary approach is fueling innovations across diverse fields, from genome editing to synthetic biology.

One key area benefiting from these innovations is epigenetic research, particularly in DNA methylation analysis. While bisulfite sequencing has traditionally been the gold standard for detecting methylated cytosines, enzyme-based deamination strategies have recently emerged as less damaging alternatives (Schutsky et al. 2018; Vaisvila et al. 2021). These enzyme-based methods better preserve input DNA integrity (Z. Sun et al. 2021), leading to improved library yields and enhanced sequencing coverage. To leverage these deaminases for methylome sequencing, complementary enzymes are often required to selectively block deamination of specific cytosine derivatives. For instance, TET enzymes, which oxidize 5mC to further derivatives like 5hmC, are integral to several methylome sequencing approaches (Y. Liu et al. 2019; Vaisvila et al. 2021). Similarly, engineered methylases have been used in protecting C from enzymatic deamination (Wang et al. 2023).

Expanding the repertoire of such deaminases and auxiliary enzymes is an area of active research as exemplified by recent reports describing novel deaminases with novel substrate specificity (Vaisvila et al. 2024; Huang et al. 2023). However, sequencing methods using these deaminases remain indirect as they rely on the deamination of canonical cytosine (C). These indirect readout methods have limitations, including a reduction in sequence complexity—effectively converting a 4-letter genetic code to a 3-letter one—and the need for complete deamination, as any residual cytosine could be misinterpreted as methylated. Consequently, there is a growing demand for more direct methods resulting from specific deamination of methylated cytosines.

In our previous work, we developed MetaGPA, a framework to link viral genotypes with specific phenotypes directly from high-throughput sequencing of environmental metaviromes (Yang et al. 2021). Using direct enzymatic treatment of nucleic acids for phenotypic selection, MetaGPA successfully identified diverse enzyme families associated with modified cytosine phenotypes, including a novel DNA hydroxymethylcytosine(5hmC) carbamoyltransferase with application to detect 5hmC in DNA (Yang et al. 2022). In this report, we apply a MetaGPA screen for cytosine modifications in metavirome DNA and identify the dCMP deaminase domain (Pfam #PF00383, dCMP_cyt_deam_1) as strongly associated with recovered sequences. We hypothesize that these deaminases are encoded by viruses containing methylated and/or hydroxymethylated cytosine fully replacing canonical cytosine in their virion DNA and are specifically adapted to the deamination of modified cytosines.

Using MetaGPA screens and sequence mining (Yang et al. 2021), we identified 284 diverse putative bacteriophage mononucleotide cytosine deaminases which were subsequently evaluated for activity on canonical and modified cytosine nucleobases in both mononucleotide and polymeric forms. Among the 56 active deaminases recovered, more than 60% exhibit pronounced selectivity towards 5mdCTP and/or 5hmdCTP over dCMP or dCTP. Furthermore, we demonstrate that these mononucleotide cytidine deaminases are active on 5mC in the context of ssDNA, supporting their application in the simultaneous sequencing of methylomes and genomes. We tested this approach in various applications, notably for the identification of rare methylation events.

## Results

### 1. MetaGPA screens identify a category of mononucleotide deaminases associated with DNA cytosine modification in bacteriophages

To identify candidate enzymes with activity towards non-canonical cytidines, we applied the MetaGPA framework (Yang et al. 2021) to a coastal seawater metavirome. For the molecular selection step, we used a cocktail of restriction endonucleases (RE) known to be blocked by cytosine modified at the C-5 position (Flodman et al. 2020). Thus, the fraction of genomic DNA that is resistant to this enzymatic treatment potentially harbors modified cytidine. To identify genetic determinants linked to this resistance, the RE digested library was amplified, sequenced, and compared to an untreated control library (**Supplementary Figure 1A**). A six frame translation followed by comparison to a Pfam-A reference database of hidden Markov model (HMM) sequence profiles identifies the domain composition in both the total metavirome library and the RE resistant fraction. Those annotated protein domains significantly enriched in contigs recovered from RE treated libraries constitute candidates for catalytic activity on modified cytosines.

Among the top five significant associations is the deaminase domain matching “MafB19-like deaminase” (PF14437) and “Cytidine and deoxycytidylate deaminase zinc-binding region” (PF00383) (**Supplementary Figure 1B**). Representatives from each of these enzyme families have previously been demonstrated to catalyze the conversion of deoxycytidine monophosphate (dCMP) to deoxyuridine monophosphate (dUMP) (Bianchi, Pontis, and Reichard 1987), a key step in the deoxythymidine biosynthetic pathway. Notably, these deaminase domains were also significantly associated with a broader definition of cytosine modifications from a previous MetaGPA screen (Yang et al. 2021), where the selection process consisted of eliminating DNA containing unmodified cytosines. We therefore combined these deaminase genes with the ones derived from the previously described MetaGPA study and identified a total of 426 genes for which 149 were from contigs predicted to contain modified cytosines. Alignment of the protein sequence reveals clustering of the annotated dCMP deaminase domains from predicted modified contigs in several branches (**Figure 1A**). 104 out of the 149 (70%) dCMP deaminase genes found in modified contigs co-localized within 6kb (+/− 3kb) of a thymidylate synthase gene. In contrast, this colocalization was rarely found (17 out of 277, 6.1%) in genomes with predicted canonical DNA (**Figure 1B, Supplementary Figure 1C**). Subsequent multiple sequence alignments of these deaminase-associated thymidylate synthase homologs revealed amino acids in their active sites that correlate with a preference for dCMP over dUMP (L. Liu and Santi 1992) (**Figure 1C**). Based on these residues, a majority (85.1%) of the 104 deaminase-associated thymidylate synthases found in modified contigs are predicted to make 5hmdC or 5mdC while the majority (52.9%) of the 17 found in unmodified contigs are predicted to make dT (Weigele and Raleigh 2016; Yang et al. 2021).

**Figure 1:**
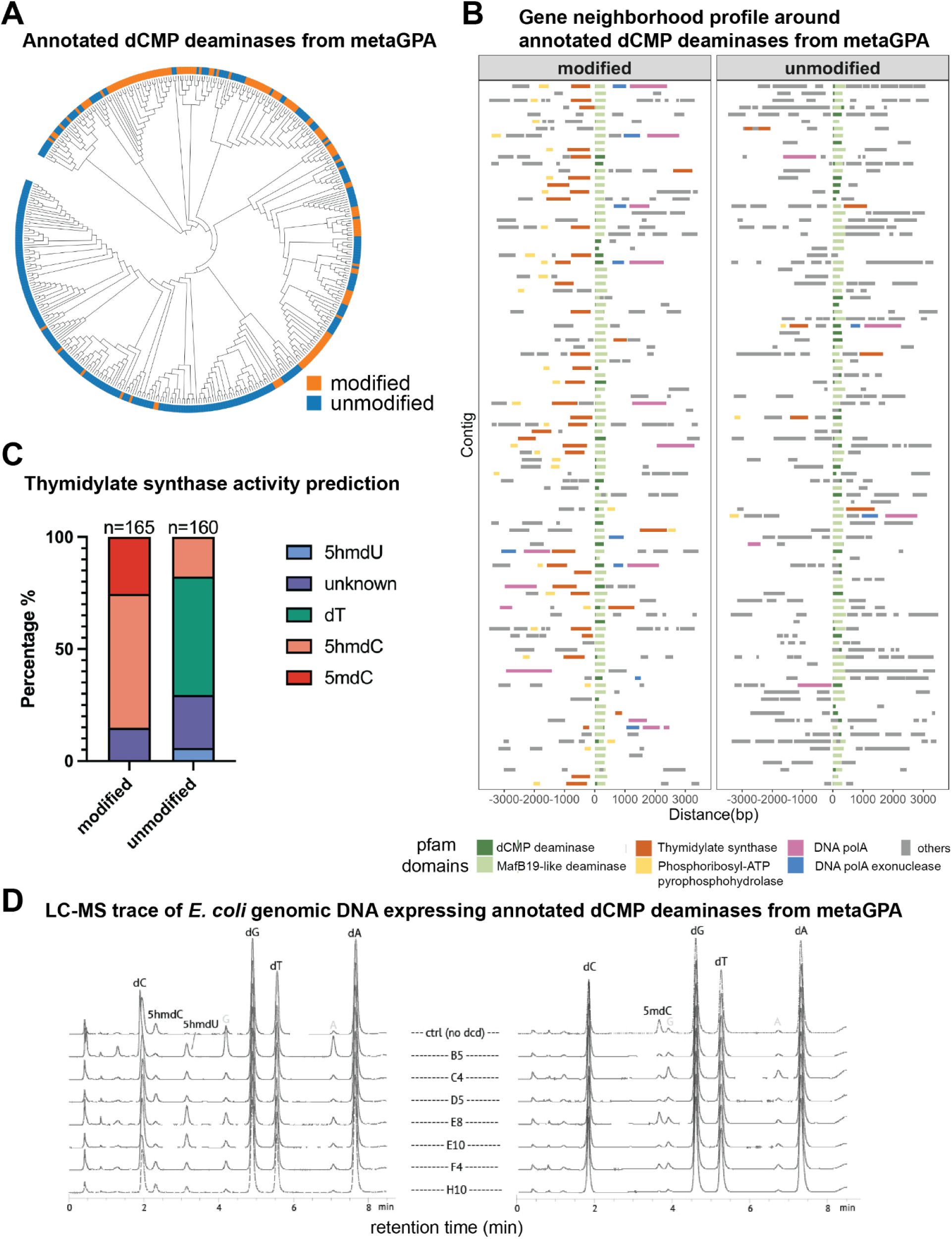
Identification of 5mC selective deaminases from MetaGPA study. **A)** Phylogenetic tree of deaminases (PFAM annotation: dCMP_cyt_deam_1, PF00383) identified from MetaGPA. The colored marks denote the source of the deaminases: orange and blue indicate dCMP Deaminases from contig predicted to have modified cytosines and canonical cytosines respectively. **B)** Gene neighborhood profiles of the dCMP deaminases within a maximum of +/− 3kb genomic region. Color bar represents the top five domains co-localized with dCMP deaminase, namely thymidylate synthase (thymidylat_synt), MafB19 like deaminase (MafB19-deam), Phosphoribosyl-ATP pyrophosphohydrolase (PRA-PH), DNA polymerase A (DNA_pol_A) and 3’-5’ exonuclease (DNA_pol_A_exo1). 104 out of 149 (70.0%) predicted modified contigs (left) containing dCMP deaminase domain also contain a thymidylate synthase domain as opposed to 17 out of 277 (6.1%) for unmodified contigs (right). For clarity only 100 randomly chosen contigs from each category are represented (for full dataset, see Supplementary Figure 1C). **C)** Predicted pyrimidine product profiles of the thymidylate synthases extracted from the MetaGPA modified (left bar) and unmodified (right bar) contig pools. dT for thymidine, 5hmdC for 5-hydroxymethyl-2’-deoxycytidine, 5hmdU for 5-hydroxymethyl-2’-deoxyuridine, or 5mdC for 5-methyl-2’-deoxycytidine). Gene neighborhood profiles of the dCMP deaminases within a maximum of +/− 3kb genomic region. Color bar represents the top five domains co-localized with dCMP deaminase, namely thymidylate synthase (thymidylat_synt), MafB19 like deaminase (MafB19-deam), Phosphoribosyl-ATP pyrophosphohydrolase (PRA-PH), DNA polymerase A (DNA_pol_A) and 3’-5’ exonuclease (DNA_pol_A_exo1). 104 out of 149 (70.0%) predicted modified contigs (left) containing dCMP deaminase domain also contain a thymidylate synthase domain as opposed to 17 out of 277 (6.1%) for unmodified contigs (right). For clarity only 100 randomly chosen contigs from each category are represented (for full dataset, see Supplementary Figure 1C). **D)** LC-MS traces of the seven annotated dCMP deaminase enzymes showing deamination activity on 5hmdC (left) and on 5mdC (right). Ctrl corresponds to control with no deaminases, B5 is mSCD-B5.

These deaminases are expected to be encoded by phages that have replaced cytosines with modified cytosines in their DNA and we hypothesized that they might deaminate C-5 modified dCMP. To experimentally validate this hypothesis, a total of 90 full length annotated dCMP deaminase genes derived from modified contigs and in cis with a thymidylate synthase were selected for an *in vivo* screen where candidate deaminases were each co-expressed with a genetic system that introduces 5-hmdCTP into the cellular dNTP pool of *E. coli* where it is subsequently incorporated *in-vivo* into DNA by the cell’s replicative DNA polymerase. The system includes bacteriophage T4 5-hydroxymethyl-2’-deoxycytidine monophosphate (5hmdCMP) synthase (gene 42) and dNMP kinase (gene 1) (**Supplementary Figure 1D**). Expression of T4 gp42 and gp1 leads to the synthesis and introduction of 5hmdCTP into the cellular dNTP pool of *E. coli* where it is subsequently incorporated *in-vivo* into DNA by the cell’s replicative DNA polymerase. If any form of 5hmdC can be a substrate for the candidate deaminase we would expect to find its deamination product, 5-hydroxymethyl-2’-deoxyuridine (5hmdU) in DNA recovered from cells in the screen. As expected, control strains not expressing a candidate deaminase show no detectable accumulation of 5hmdU in recovered plasmid DNA subjected to nucleoside analysis by LC-MS. For seven of the candidates expressed under these conditions, we observed an accumulation of 5hmdU (confirmed mass of 258.2 u by MS) in recovered plasmid DNA (**Figure 1D**). We interpret these data as these seven candidates deaminating 5hmdC at some step(s) in the biosynthetic pathway from 5hmdCMP to DNA.

Based on the identity of the residue at the position homologous to N177 of E. coli ThyA, the thymidylate synthase of all seven deaminases are predicted *in-silico* to convert dCMP to 5mdCMP (**Supplementary Figure 1E**). To confirm our *in-silico* predictions, we experimentally tested the activity of these thymidylate synthases. For this, the annotated open reading frame was cloned and expressed in *E. coli* together with bacteriophage T4 gp1, a promiscuous dNMP kinase capable of phosphorylating non-canonical pyrimidines en route to the dNTP pool. Following expression of these two enzymes, plasmid DNA was recovered from cells following expression of these two enzymes and analyzed by LC-MS. The thymidylate synthase of the *Xanthomonas* phage Xp12, which is characterized by the complete substitution of cytosine with 5-methylcytosine (5mC) in its genomic DNA (Farber and Ehrlich 1980), was included as a positive control. Five out of the seven thymidylate synthases produced 5mdCMP (**Supplementary Figure 1E**), suggesting that the associated in-cis deaminases are involved in the biosynthesis of dTMP from 5mdCMP.

Of the seven deaminases selected for further characterization, one candidate termed B5, demonstrated good solubility after recombinant cloning and expression and showed the highest yield of 5hmdU in the *in-vivo* assay. The thymidylate synthase linked to this deaminase catalyzes the production of 5mdCMP (**Supplementary Figure 1E**), with homologs identified in bacteriophages like Xanthomonas phage Xp12 (Kuo and Tu 1976) and Pseudomonas phage PaMx74 (**Supplementary Figure 2A**), which replace C entirely with 5mC in their genome. This suggests a similar substitution pattern in the B5 phage. B5 was therefore hypothesized to be a Methylation-Selective Cytidine Deaminase (mSCD) and designated “*mSCD-B5*” for further study.

### 2. mSCD-B5 selectively deaminates 5mC and 5hmC in nucleoside triphosphate and polynucleotide forms

mSCD-B5 gene is originally from a contig assembled from the metagenomic sequence of the viral fraction in a wastewater treatment plant microbiome (Yang et al. 2021). mSCD-B5 is a single domain protein, consisting of 154 amino acids with a predicted molecular mass of 17.04 kDa (**Supplementary Figure 2B**). An AlphaFold2 structural model reveals a prototypical mononucleotide deoxycytidine deaminase fold (**Supplementary Figure 2C**). Its closest structural homolog in the PDB is a cytidine deaminase from the *Chlorella* virus PBCV-1 (Y.-H. Li et al. 2022)(Zhang et al. 2007) (**Supplementary Figure 2C**). Other structurally similar proteins include the cytidine deaminases of cyanophage S-TIM5 (Marx and Alian 2015) and *Escherichia* bacteriophage T4.

To characterize the biochemical activities of mSCD-B5, the enzyme was recombinantly expressed and purified (See **Materials and Method**, **Supplementary Figure 2B**). Deamination reactions were performed with purified enzymes using various cytosine nucleotide substrates, and reactions were monitored and quantified by LC-MS. mSCD-B5 is an active cytosine deaminase, with a marked preference for cytosine methylated at C-5 position (5mC). Selective deamination activities, particularly on 5mC but not on C nucleotide triphosphates were confirmed with LC-MS. By thirty minutes, ninety-nine percent of the 5mdCTP was deaminated by mSCD-B5, contrasting with 4% of dCTP deaminated over the same reaction time (**Figure 2A-B**). mSCD-B5 also efficiently deaminated 5hmdCTP, albeit at a slower rate than 5mdCTP (65% over 30 minutes). Additionally, the enzyme showed a clear preference for nucleotide triphosphate versus monophosphates with no detectable activity observed on either dCMP or 5mdCMP (**Figure 2A-B**).

**Figure 2:**
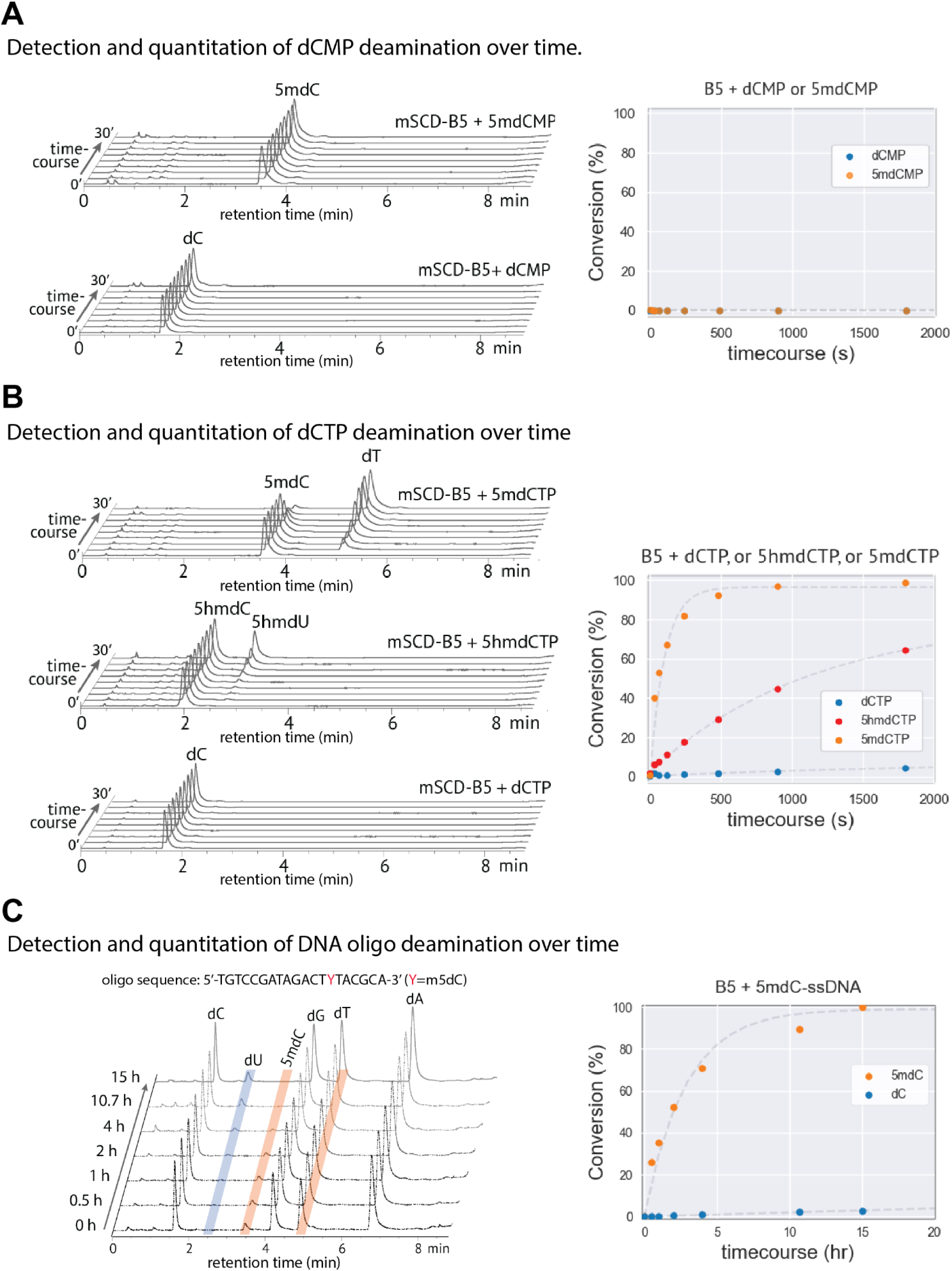
mSCD-B5 deamination rate *in vitro*. **A)** Time courses of deamination reactions with mononucleotide monophosphate substrates analyzed by LC-MS (5mdCMP, top traces and dCMP, bottom traces). Time courses range from 0 to 30 minutes of mSCD-B5 treatment. Left : raw LC-MS traces; Right : graphical representation of the conversion rate (in percent) of 5mdCMP to dTMP (orange) and dCMP to dUMP (blue) function of time (in second). **B)** Time courses of deamination reactions with mononucleotide triphosphate substrates analyzed by LC-MS (5mdCTP, top traces, 5hmdCTP, middle traces and dCTP, bottom traces). Time courses range from 0 to 30 minutes of mSCD-B5 treatment. Left : raw LC-MS traces; Right : graphical representation of the conversion rate (in percent) of 5mdCTP to dTTP (orange), 5hmdCTP to 5hmdUTP (red) and dCTP to dUTP (blue) function of time (in second). 250 µM nucleotide phosphates and 5 µM mSCD-B5 were used per reaction. **C)** Time courses of deamination reactions with oligonucleotide substrates containing either dC or 5mdC analzyed by LC-MS. Time courses range from 0 to 15 hours of mSCD-B5 treatment. Left : raw LC-MS traces; Right : graphical representation of the conversion rate (in percent) of 5mdC to dT (orange) and dCto dU (blue) function of time (in hours). For all graphical representations, the conversion rates (in %) were calculated from the integration of the pick area of both unconverted and converted base as measured by LC-MS.

Given the robust *in vitro* activity of mSCD-B5 on triphosphorylated nucleotides, and previous reports suggesting that mononucleotide enzymes can be directed to act on polynucleotides, we investigated whether mSCD-B5 could deaminate 5mC on DNA. We performed deamination reactions on single-stranded oligonucleotides carrying internal 5mC or 5hmC and analyzed the appearance of deamination products using LC-MS (**Figure 2C, Supplementary Figure 2D**). Similar to what we observed with mononucleotide substrates, mSCD-B5 selectively deaminates 5mC or 5hmC on single-stranded polynucleotide substrates with minimal activity on C.

We next assessed the activity of mSCD-B5 on double-stranded DNA (dsDNA). To do this, mSCD-B5 deamination reactions were performed directly on, or following heat denaturation of, dsDNA from bacteriophages containing 5mC (Xp12), 5hmC (T4*gt*, a mutant of T4 deficient in DNA α- and β-glucosyltransferases and as a result contains 5hmC in its genome (Miller et al. 2003)), or unmodified C (Lambda). Using LC-MS, we observed minimal deaminase activity on intact dsDNA from all bacteriophages, while heat-denatured (ssDNA) samples showed substantial deamination, with 35% of 5mC and 27% of 5hmC converted (**Supplementary Figure 2E**). These findings indicate that mSCD-B5 deaminase activity can occur on complex ssDNA samples produced by denaturation, but is either abolished or greatly reduced on native dsDNA.

Taken together, our data demonstrate that mSCD-B5 is a deoxycytidylate deaminase with methylation selective cytidine deamination activity on ssDNA *in-vitro*.

### 3. Substrate range of mSCD-B5 deaminase

Having established mSCD-B5’s preferential activity on 5mC and 5hmC in the context of single-stranded DNA (ssDNA), we further investigated its substrate range and sequence specificity using a combination of high-throughput sequencing and analytical methods.

#### Deaminase activity on all sequence contexts and positions in the ssDNA fragment

To determine whether mSCD-B5 deamination occurs preferentially within specific sequence contexts or positions on the ssDNA molecule, we performed high throughput short-read sequencing on a *gDNA-mix* library treated with mSCD-B5 (**Figure 3A, Materials and Methods**). The *gDNA-mix* library is composed of genomic DNA from bacteriophages Xp12, T4gt, and Lambda, which represent fully 5mC-modified, fully 5hmC-modified, and unmodified cytosine (C) genomes, respectively. Additionally, we incorporated genomic DNA from *E. coli* K12 strain DHB4 (dcm+), which harbors 5mC in the C<u>C</u>WGG context (where W = A or T, with methylation occurring on the internal cytosine) and a pUC19-plasmid for which 5mC methylation is restricted to CpG context. Both *E. coli* and pUC19 provide DNA with mixed cytosine modification states with pUC19 mimicking human genome methylation.

**Figure 3:**
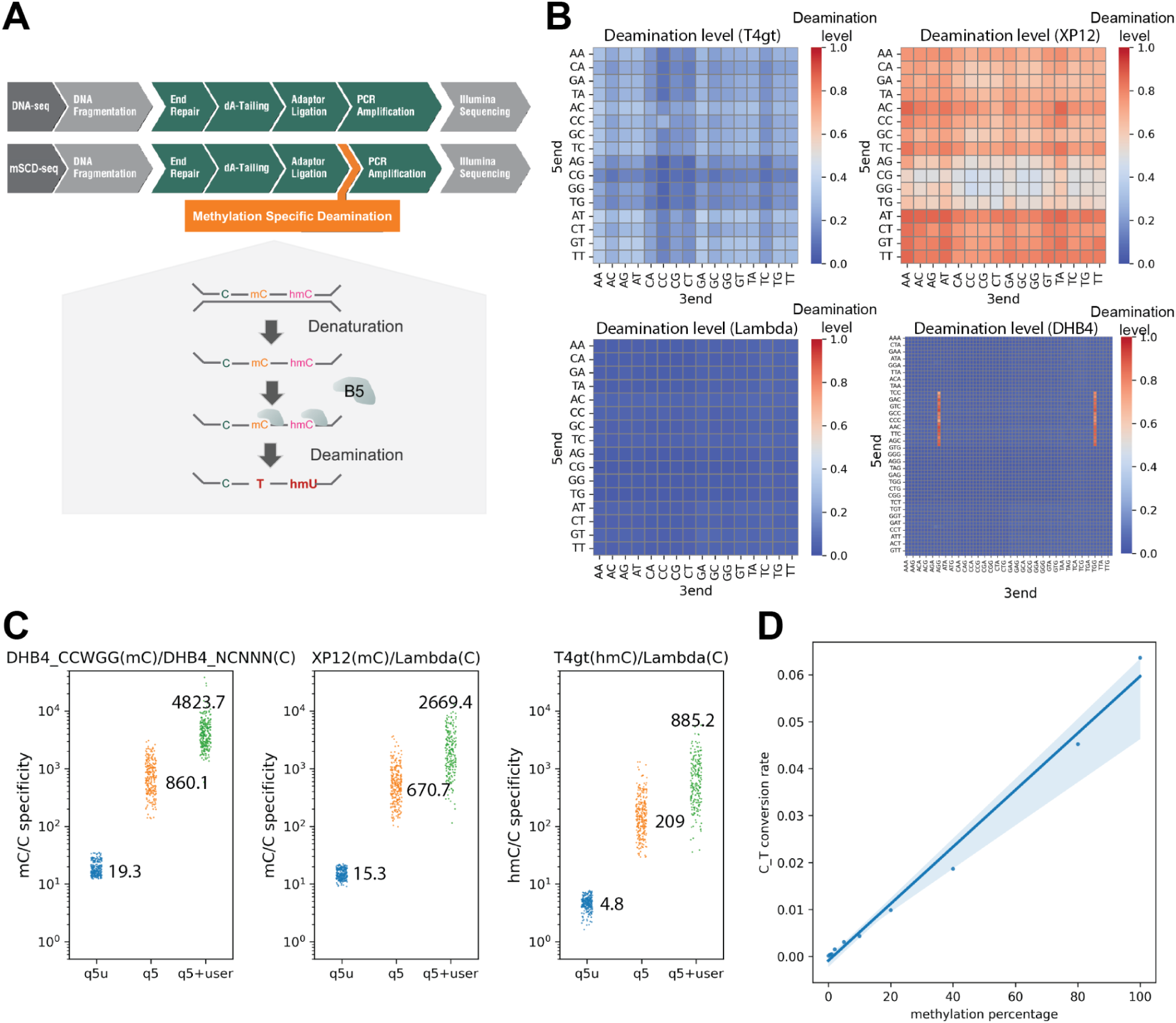
High-throughput sequencing of mSCD-B5 treated libraries. **A)** Workflow comparison of short-read sequencing using mSCD-B5 versus conventional DNA-seq. The denaturation and deamination steps are integrated into the standard library preparation protocol after adaptor ligation. The remaining steps follow standard DNA-seq library preparation without requiring conversion-resistant adapters or dU-compatible polymerases. **B**) Heatmaps of the average deamination level (C-to-T conversion rate) in all NNCNN sequence contexts (in T4gt (5hmC), XP12(5mC), Lamda (C)) or all NNNCNNN sequence context (in dcm+ *E. coli* DHB4). Color scale ranges from 0.0 to 1.0 (0 to 100% deamination) **C**) Stripplots showing ratio between the C-to-T conversion rates in C<u>C</u>WGG compared to all other NCNNN contexts (excluding C<u>C</u>WGG, in E.coli DHB4, left plot); the ratio between C-to-T conversion rates in XP12 (5mC) compared to Lambda (C) in all NNCNN contexts (middle plot) and the ratio between C-to-T conversion rates in T4gt (5hmC) compared to Lambda (C) in all NNCNN contexts (Right plot) when mSCD-B5 treated libraries were amplified with Q5U (blue), Q5 (orange) and treated with USER and amplified by Q5 (green). Numbers in plot represent average specificity ratio. **D**) Regression plot showing relationship of methylation levels (in synthetic oligo, x axis) and detected deamination level (y axis).

In brief, fragmented genomic DNA was ligated with standard sequencing adapters and the resulting libraries were denatured and treated with mSCD-B5 deaminase at 42 °C for one hour prior to amplification and next-generation sequencing (NGS) (**Supplementary Figure 3A-E, Supplementary text 1** and **Materials and Methods**). In order to capture all deamination events including deamination of canonical C, amplification was performed using the uracil tolerant engineered polymerase Q5U (Wardle et al. 2008). Because deamination is preferential for 5mC sites, reads originated from sparsely methylated or unmethylated genomes such as *E. coli*, pUC19 and Lambda control have mostly retained the same DNA sequence and can be mapped using standard mapping algorithms such as Bowtie2 or BWA-MEM. Conversely, reads originated from Xp12 and T4gt are expected to be largely converted and require conversion-aware mapper such as Bismap or BWA-Meth. Comparison of all these tools indicate that the standard BWA-MEM mapper provided the best methylation calls in *E.coli* (**Supplementary text1**, **Supplementary Figure 4A**). Standard quality control metrics for NGS reveal size distribution and GC bias that are in line with those obtained from standard DNA-seq indicating that mSCD-B5 treatment followed by Q5U amplification does not significantly degrade DNA (**Supplementary Figure 4C-D**)

To evaluate deamination in specific sequence context, the deamination rates at each genomes were calculated for each of the 256 NN<u>C</u>NN or N<u>C</u>NNN contexts (where N=A,T,C or G). Consistent with our LC-MS observations on ssDNA oligos and denatured genomic DNA, we observe significant deamination levels for 5mC and 5hmC (**Figure 3B**). We show that the deamination yield on 5mC (in Xp12) ranges from 44.4% to 87.5% with a mean value of 69.3% depending on the NN5m<u>C</u>NN sequence context (**Supplementary Figure 4B**). Noticeably, the deamination yields at the respective NNCpGN contexts in pUC19 ranged from 49.1% to 95.5% which are on average 10% higher than those in Xp12, suggesting that fully methylated DNA presents a more challenging substrate than sparsely methylated DNA (**Supplementary Figure 4B**). Using T4gt, we evaluated the deamination of 5hmC to range from 4.8% to 39.1% with a mean value of 22.4% respectively (**Supplementary Figure 4B**). On *E. coli* genomic DNA, mSCD-B5 selectively deaminates mC with an average C-to-T conversion of 84.7% in C<u>C</u>WGG context while non-specific deamination at C with an average deamination rate of 3.7%, representing a nearly 20-fold specificity ratio for deamination of 5mC over C by mSCD-B5 (**Figure 3C**).

Finally, to assess the relationship between deamination and methylation levels, we varied the ratio of 5mC:C in a dilution series using an oligonucleotide carrying internal, unmodified C mixed with an oligonucleotide identical in sequence but carrying an internal, methylated C. High throughput sequencing of the libraries treated with mSCD-B5 reveals a linear relationship between the methylation percentage and deamination indicating a quantitative measurement of methylation levels by mSCD-B5 (**Figure 3D, Supplementary Figure 4E**).

#### Cytosine deamination by mSCD-B5 is restricted to certain derivatives modified at position 5

In addition to 5mC, prokaryotes often contain 4mC and 6mA at specific sites in their genomic DNA (Blow et al. 2016). To evaluate the activity of mSCD-B5 on common prokaryotic DNA modifications, we performed NGS on a mSCD-B5-treated library consisting of a gDNA-mix and genomic DNA from *Natrinema pallidum* BOL6-1 (*N. pallidum*). This strain is known to express both 6mA and 4mC methyltransferases, producing the methylated motifs 4mCTAG, C6mATTC, GTAYT4mCG, and CAGYA6mAC (DasSarma et al. 2019). To precisely measure deamination levels, the fraction of C-to-T or A-to-G conversions in the second paired read was subtracted from that in the first paired read in order to specifically measure damage as opposed to sequencing artifact and variation (Chen et al. 2017)

We observed that deamination levels at 4mC-methylated <u>C</u>TAG (**Supplementary Figure 5A,C**) and GTAYT<u>C</u>G contexts in *N. pallidum* (**Supplementary Figure 5B**) are nearly at the level of sequencing error, being at least ten times lower than in unmethylated contexts. In contrast, unmethylated CTAG and GTAYTCG contexts from lambda genome exhibit deamination levels comparable to other contexts. Taken together, these results indicate that mSCD-B5 does not deaminate cytosines containing a methyl group at position 4. To investigate whether mSCD-B5 is capable of deaminating A or 6mA, we examined the excess of A-to-G transitions in all contexts including 6mA-modified contexts. We observed that the excess of A-to-G transition frequency at C<u>A</u>TTC (∼0.00003) in *N. pallidum* was within the levels of standard sequencing error rates (**Supplementary Figure 5D**), indicating that mSCD-B5 lacks deaminase activity toward adenine or its derivative, 6mA. Overall, when applied to prokaryotic genomic DNA—where 6mA, 4mC, and 5mC are the predominant modifications— mSCD-B5 selectively identifies 5mC modifications, as neither 4mC nor 6mA are substrates for deamination.

#### Oxidized and glucosylated products of 5hmC are not substrates of mSCD-B5

Finally, we tested single stranded oligonucleotides carrying internal 5-formylC (5fC), 5-carboxyC (5caC) and 5-glucosylmethylcytosine (5gmC) and saw no detectable activity of mSCD-B5 on these substrates using LC-MS readout (**Supplementary Figure 2D**). We also saw no detectable activity of mSCD-B5 deaminases on denatured T4 genomic DNA indicating that both α- and β-5gmC are resistant to deamination by mSCD-B5 (**Supplementary Figure 2E**).

### 4. mSCD-B5 activity combined with BGT provides 5mC specific readout in methylome sequencing

Having established that α- and β-5gmC are not substrates of mSCD-B5 (**Supplementary Figure 2D-E**), we sought to identify methylated sites in DNA containing a mixture of 5mC and 5hmC. The treatment of 5hmC with T4 phage β-glucosyltransferase (BGT) producing β-5-gmC has been previously used to protect β-5gmC from deamination by APOBEC (Z. Sun et al. 2021). Therefore, we used high-throughput sequencing to investigate whether BGT treatment prior to deamination could achieve a 5mC specific readout by fully protecting 5hmC from mSCD-B5-mediated deamination.

To do so, the *gDNA-mix* was treated with or without BGT prior to deamination (**Materials and Methods**). The resulting libraries were amplified with Q5U to capture all deamination events and sequenced to an average of 6.7 million paired-end reads. Reads were mapped to the reference genomes using BWA-Meth and analyzed for C-to-T imbalance between Read 1 and Read 2, a sensitive measurement for detecting deamination events. The method was previously shown to distinguish deamination from sequencing errors or variants (L. Chen et al. 2017) and to be a particularly sensitive measure for low levels of deamination.

Consistent with mSCD-B5 activity, the control sample without BGT treatment shows significant deamination on both 5mC and 5hmC, while unmodified C exhibits minimal deamination (<0.5%) (**Figure 4A**). BGT treatment does not alter the deamination rates of both unmodified C and 5mC but entirely suppresses deamination of 5hmC (<0.1%) across all sequence contexts (**Figure 4A-B**). This result confirms that β-glucosyl-methylcytosine is not a substrate for mSCD-B5, and pre-treating DNA with BGT entirely blocks deamination at 5hmC sites. Combined with our LC-MS assay showing that 5-formylcytosine (5fC) and 5-carboxylcytosine (5caC) are also not substrates for mSCD-B5 (**Supplementary Figure 2D**), we conclude that BGT treatment prior to mSCD-B5 deamination would selectively reveal 5mC in genomes (such as the human genome) containing a mixture of C, 5mC, 5hmC, 5fC, and 5caC.

**Figure 4:**
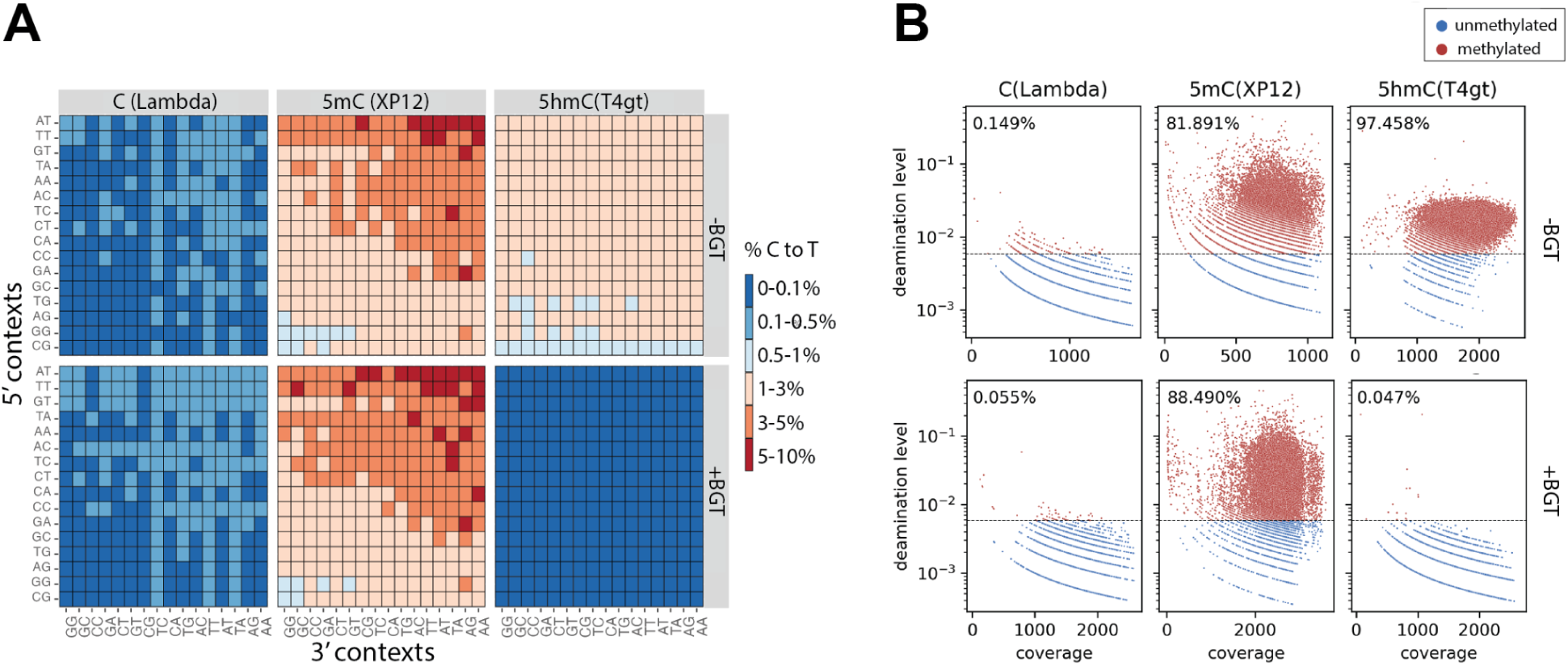
mSCD-B5 with prior BGT treatment provides 5mC specific detection. **A)** Heatmap plots showing deamination level for all NNCNN contexts with (bottom panels) or without (top panels) BGT treatment in Lambda (left), XP12 (middle) and T4gt (right) **B**) Scatter plot showing deamination levels at base resolution function of read coverage with (bottom panels) or without (top panels) BGT treatment. Methylation threshold is displayed as a dotted line in each plot. The number in each plot represents the percentage of genomic sites that are above the threshold in Lambda (left), XP12 (middle) and T4gt (right).

### 5. USER Treatment Enhances mSCD Specificity for 5mC compared to C by >4000-Fold

Since the deamination of canonical C, 5mC, and 5hmC produces U, T and and 5-hydroxymethyluracil (hmdU) respectively, it is possible to selectively eliminate the background deamination of canonical C. This can be achieved by using DNA polymerases containing a uracil recognition pocket, such as those found in archaeal family B DNA polymerases (Connolly 2009), commonly used for NGS library amplification. Alternatively or in combination, residual dU can be removed prior to amplification using combination of Uracil DNA Glycosylase (UDG) (Lindahl et al. 1977) and DNA glycosylase-lyase Endonuclease VIII, also known as USER treatment (Bitinaite et al. 2007). To demonstrate the elimination of dU resulting from canonical C deamination, mSCD-B5-treated *gDNA-mix* libraries were either amplified with Q5U or Q5 with or without prior treatment with USER and sequenced. On library performances, we observed a loss of material (ranging from 4-8 fold decrease) in Q5 and Q5+USER amplification compared to Q5U. Accordingly, quality control metrics show biases in base composition, with an overrepresentation of AT-rich regions and smaller insert sizes observed in libraries treated with Q5 or USER/Q5 (**Supplementary Figure 4D**). This result is in agreement with the elimination by USER/Q5 of fragments containing dU as longer and GC-rich fragments are preferentially eliminated.

Regarding the deamination ratio, we observed an average of 19 fold difference between deamination of 5mC compared to C for sample amplified using Q5U. This difference increased to an average of 860 fold when the sample is amplified with Q5 instead of Q5U and to an average of 4823-fold when the sample is treated with USER prior to amplification with Q5. Similarly, amplification using Q5U, Q5 and USER+Q5 leads to an average 5 fold, 209 fold and 885 fold in apparent deamination levels of 5hmC compared to C respectively (**Figure 3C**). These results indicate that the selectivity for 5mC/5hmC can be significantly enhanced by applying a USER treatment to the deamination reaction. The highest specificity is achieved when combining post-deamination USER treatment with Q5 polymerase. Under these conditions, the absolute C-to-T conversion rate at unmethylated cytosine (in phage lambda genome) reaches values as low as 0.02% in average (ranging from 0% to 0.19%) (**Supplementary Figure 4B**), which closely approaches, if not matches, the baseline error rate of Illumina sequencing. With these optimized conditions, the mSCD-B5 deamination step followed by USER treatment offers unparalleled specificity for detecting 5mC/5hmC.

### 6. Direct conversion of 5mC by mSCD-B5 identifies methylation at unprecedented sensitivity

This unparalleled specificity, achieved by completely eliminating background deamination of unmodified cytosine, offers a clear advantage in enhancing the sensitivity of methylome sequencing. To demonstrate its utility, we refined experimental protocols and data analysis pipelines to capture and leverage this sensitivity across various test cases :

#### Partial methylation of control lambda DNA in C<u>C</u>WGG context

Sequencing the *gDNA-mix* library treated with mSCD-B5 followed by USER+Q5 treatment, we observed residual methylation levels in an ostensibly unmethylated lambda control at C<u>C</u>WGG sites (**Supplementary Figure 6A**). Unmethylated lambda genomic DNA – a commonly employed control for unmethylated DNA in methylome sequencing – is typically derived by propagation of the phage in a *dcm^-^ E.coli* host; a mutant strain bearing the *dcm-6* allele, which contains a nonsense mutation in the *dcm* gene (Dar and Bhagwat 1993). Thus, we speculate that *dcm*-6 strains retain residual *dcm* methyltransferase activity. To demonstrate the presence of residual methylation at C<u>C</u>WGG sites in *dcm*-6 strains, libraries of *E.coli* strains containing either wild-type *dcm* (*dcm^+^*) strain DHB4, a nonsense mutation strain ER3662 in the dcm gene (*dcm*-6), or a complete deletion the *dcm* gene (*Δdcm,* strain ER2861) were treated with mSCD-B5 followed by USER treatment. We observed a strong and weak methylation at C<u>C</u>WGG for the *dcm^+^* and *dcm-6* strains, respectively, and no detectable methylation in the *Δdcm* strain (**Supplementary Figure 6B**). We validated this result using Tet-assisted PacBio sequencing (Clark et al. 2013) (**Supplementary Figure 6C**).

This experiment demonstrates the high sensitivity of 5mC detection when combining mSCD-B5 deamination with subsequent USER treatment prior to amplification with Q5. Notably, the detection of residual methylation in the supposedly methylation-free lambda genomic DNA emphasizes sensitivity in identifying even minute amounts of methylation.

#### mSCD-B5 deaminase provides a 0.1% methylation detection limit in standard NGS

To better assess the sensitivity of methylation detection, we prepared duplicate mixtures of genomic DNA from the *dcm^+^ E. coli* strain DHB4 and the *Δdcm* strain ER2861 (**Figure 5A**) in ratios ranging from 0.03% to 4% *dcm^+^*DNA, alongside controls including pUC19 (5mCpG), Xp12 (5mC), T4*gt* (5hmC), and lambda (unmodified C) DNA. Additionally, pure *dcm^+^* and *Δdcm* libraries were designed to provide a calibration for fully methylated sites at C5mCWGG (100%) and unmethylated sites at CCWGG (0%), respectively. The resulting mSCD-B5-treated libraries were amplified using Q5 after USER treatment to achieve the highest level of specificity. Quantitative LC-MS was performed on the 100% and 4% *dcm*+ DNA sample to provide an independent measurement of methylation (**Materials and Methods**).

**Figure 5:**
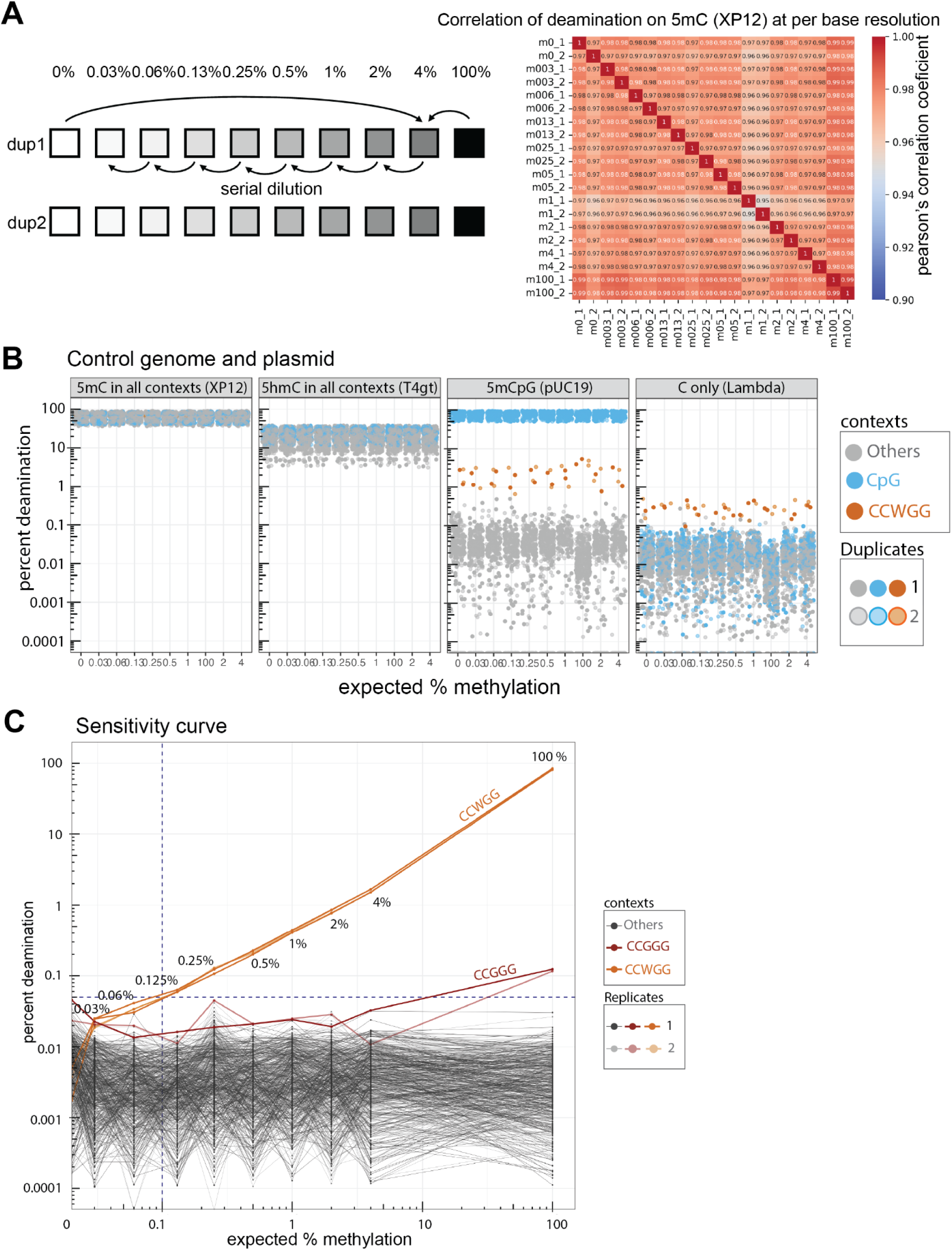
Deamination by mSCD-B5 enables detection of methylation levels as low as 0.1%. **A)** Preparation of a 2 fold serial dilution series : A sample containing 4% methylation at dcm sites was obtained by mixing *dcm+* genomic DNA (100% methylation at dcm sites) with *Δdcm* genomic DNA (0% methylation) at a 4:96 molar ratio. Serial 1:2 dilutions were performed by mixing an equimolar amount of *Δdcm* genomic DNA to the previous dilution, starting from the 4% sample. The serial dilutions generated samples containing methylation levels at 2%, 1%, 0.5%, 0.25%, 0.13%, 0.06% and 0.03% respectively Each dilution level sample was split into two duplicates. The correlation of deamination in 5mC (XP12) on per base resolution between individual samples is displayed as heatmap in the right panel. **B**) Percentage of deamination across all 256 NCNNN contexts (where N=A, T, C, or G; log10 scale on y-axis) measured on control genomic DNA samples: XP12 (5mC, left panel), T4gt (5hmC, center-left panel), pUC19 (CpG-context 5mC, center-right panel), and Lambda (canonical C, right panel) for the 10 samples analyzed. Deamination in <u>C</u>pG sites (blue) and C<u>C</u>WGG motifs (orange, where W=A or T) is highlighted to denote CpG methylation in pUC19 and the recognition sites of the Dcm methylase, respectively. **C**) Sensitivity curve representing the percentage of deamination across all 256 NCNNN contexts in E. coli DNA mixtures of Δdcm and dcm+ strains. A value of 100% methylation corresponds to DNA from a pure dcm+ strain, while 0% represents DNA from a pure Δdcm strain. Deamination in C<u>C</u>WGG (orange) and CCGGG (red) contexts is highlighted to indicate the Dcm methylase recognition sites and potential “star” activity of the Dcm methylase, respectively.

Paired-end reads were aligned to the composite genomes using BWA-MEM (for pUC19, lambda, and *E. coli* data analysis) and BWA-Meth (for Xp12 and T4gt data analysis). To accurately measure low deamination rates, the fraction of C-to-T conversions in the second read was subtracted from that in the first read of paired read (**Material and Methods**). The average deamination rates at NCNNN positions were as follows: 67.2% for 5mC (Xp12, average all contexts), 22% for 5hmC (T4gt, all contexts), 76.2% for N5mCpGNN (pUC19), and 82.2% for C5mCWGG (from *dcm^+^* E. coli) (**Figure 5B**, **Supplementary Figure 7**). By contrast, the lambda genome exhibited a low deamination rate (averaging 0.008%), reflecting minimal background activity at unmodified cytosine. Using the *dcm^+^*/*Δdcm* dilution series, we reliably detected residual methylation in C<u>C</u>WGG contexts down to 0.1% (**Figure 5C**). At this methylation level, the *dcm^+^ E. coli* strain showed residual methylation at C<u>C</u>GGG sites, suggesting star activity of the Dcm methyltransferase. Additionally, partial methylation of control lambda and pUC19 DNA were observed at C<u>C</u>WGG sites, consistent with the genetic background of their *dcm-6* host (**Figure 5B)**.

Taken together, these results demonstrate the high sensitivity of mSCD-B5 in detecting methylation, with reliable identification down to 0.1%. This sensitivity level reveals previously undetected methylation in standard controls used for methylation analysis.

#### Applications to high throughput methylome and genome sequencing in human

Most applications for detecting low-level residual methylation are focused on human studies, prompting us to apply mSCD-B5 for human methylome sequencing. To first validate that mSCD-B5 can be used for methylome sequencing in human, we sequenced mSCD-B5 treated libraries from various input of Human NA12878 genomic DNA, a well-characterized sample with numerous reference sequencing and methylation datasets available from methods such as EM-seq (Vaisvila et al. 2021), bisulfite sequencing (Ulahannan and Greally 2015), and Nanopore sequencing (Simpson et al. 2017; Rand et al. 2017).

From a direct methylome sequencing for which only 5(h)mC is deaminated (**Figure 6A, Supplementary Figure 8A)**, we expect the sequencing accuracy of canonical bases, which comprise over 99% of the human genome, to be consistent with that of standard DNA-seq. To confirm this, we analyzed all substitution frequencies in mSCD-B5-treated libraries amplified with Q5 post-USER treatment and compared these frequencies to those obtained with a “platinum” standard PCR-free DNA-seq (see **Supplementary Figure 8B, Supplementary Text1**). We observed that, beside the <u>C</u>pG context for which the C-to-T transition is elevated to an average of 60% as a result of deamination of 5(h)mC, the frequency of other substitutions, including other substitution in CpG context (C to A and C to G), are comparable to those obtained with “platinum” standard PCR-free DNA-seq. No notable GC coverage bias across the genome was observed (**Supplementary Figure 8C**). This result indicates that sequencing accuracy of NA12878 genomic DNA is comparable to standard Illumina DNA-seq, excluding C-to-T in CpG sites.

**Figure 6:**
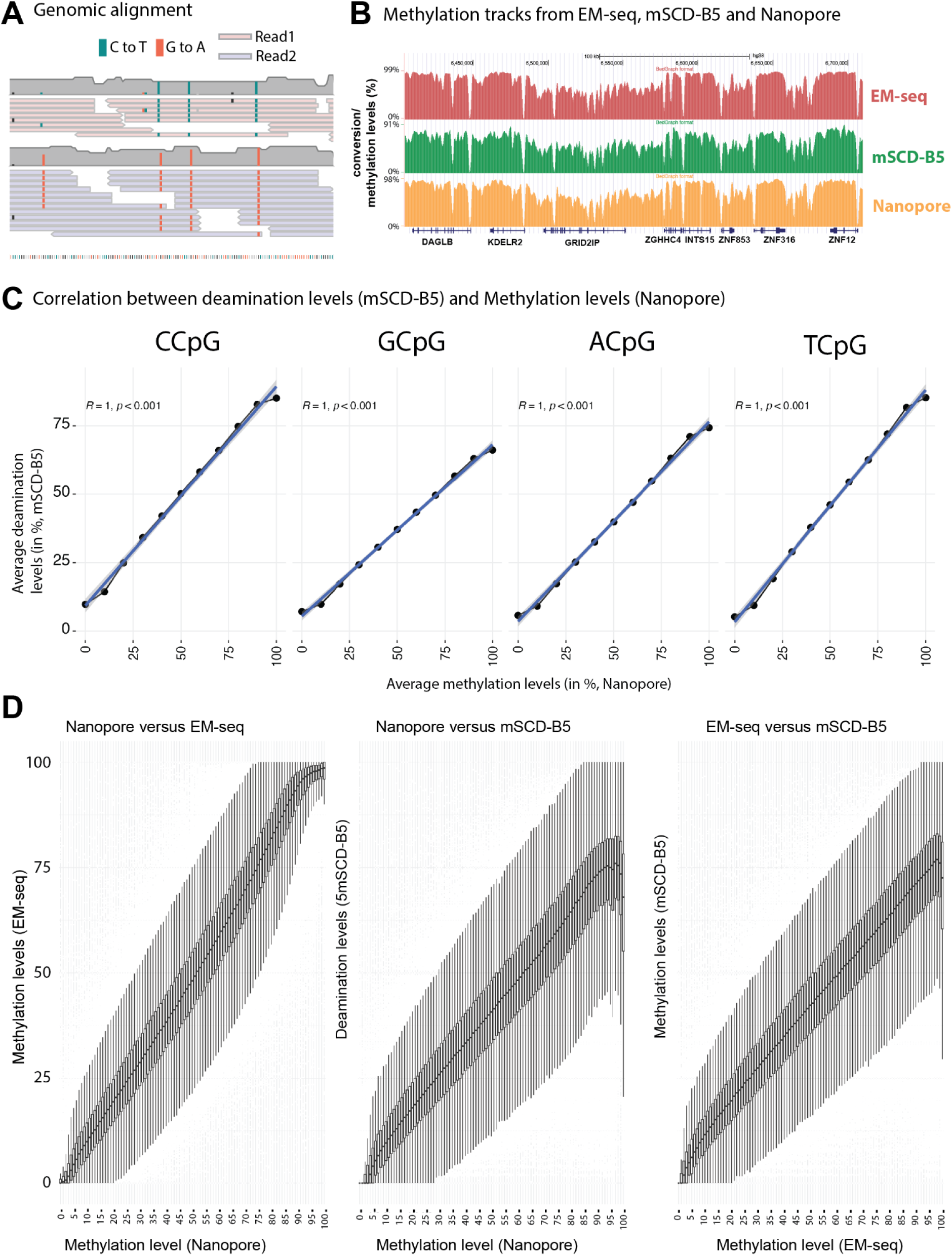
Methylome and genome sequencing of human genomic DNA from NA12878 cell line. **A)** Aligned sequencing reads (after mSCD-B5 treatment) to an example locus in the human genome. Reads were divided between paired-end read 1 (top, pink) and paired-end read 1 (bottom, blue). C-to-T transitions (green) and G-to-A transition (red) from mSCD-B5 deamination are restricted to CpG contexts and are identifying methylated sites. **B)** UCSC genome browser visualization of NA12878 methylation track from EM-seq (red) mSCD-B5 (green) and Nanopore (yellow) **C)** Genome-wide methylation level correlation between Nanopore and mSCD-B5 (starting genomic material of 100 ng). Nanopore base resolution methylation levels were binned into 0-10%, 10-20%. 90-100% (x-axis) and the equivalent methylation from mSCD-B5 were computed and plotted (Y axis) according to the NCpG context (with N=A,T,C or G). **D)** Correlation between regional methylation levels between Nanopore -EM-seq (R=0.95, p < 2.2e-16, left panel) Nanopore - mCSD-B5 (R = 0.88, p < 2.2e-16, center panel) and EM-seq - mCSD-B5 (R= 0.91, p < 2.2e-16, right panel).

Methylated positions were quantified at single-base resolution using Methyldackel (**Supplementary Text 1**). Manual visualization of 5mC levels obtained using mSCD-B5 are in good agreement with 5mC levels obtained using EM-seq and Nanopore sequencing (**Figure 6B)** and genome-wide averaged methylation levels correlated strongly with Nanopore sequencing data across all NCpG contexts (correlation = 1, p-value < 0.01, **Figure 6C**). Similar correlations were also observed for varying input DNA amounts, with detection from as little as 0.05 ng (**Supplementary Figure 8D**). However, quality control metrics and duplication rates for low input samples indicated substantial material loss during USER treatment, underscoring limitations for reduced starting DNA amounts (**Supplementary Figure 8D**). A refined comparison using sliding windows demonstrates strong concordance between methylation levels derived from EM-seq, Nanopore, and mSCD-B5-based sequencing (**Figure 6D**). Collectively, these results establish that mSCD-B5 enables simultaneous high-resolution methylome and genome sequencing.

#### Identification of low amounts of methylation in a mixed human genomic DNA

Having established that mSCD-B5 derived methylation levels are in agreement with standard methylome sequencing technologies, we sought to address the sensitivity of mSCD-B5 to identify low amounts of methylation. For this, we mixed genomic DNA from the NA12878 cell line with genomic DNA from a DNMT1 and DNMT3b double knockout DKO HCT116 cell line (Rhee et al. 2002) at a 99:1 ratio respectively. Hypomethylated CpG islands in NA12878 were selected and ranked according to the methylation levels in WT-DKO (0-30%, 30-50%, 50-80% and >80%). We can observe a gradual increase in methylation levels in the mixed sample to the target 1% in CpG islands where the WT-DKO levels are > 80% (**Supplementary Figure 8E**).

#### Application to mapping methylation in brain genomic DNA

Brain genomic DNA is characterized by non-CpG methylation and substantial levels of 5hmC. To selectively target 5mC without interference from oxidized products (5hmC, 5fC and 5caC), we applied mSCD-B5 with BGT treatment, followed by USER and Q5 amplification, to 100 ng of human brain genomic DNA spiked in with the *gDNA-mix* (**Supplementary text 1, Supplementary Figure 9A-C**). This approach enabled precise and direct base-resolution mapping of 5mC levels across CpG and non-CpG sites (**Figure 7A**). In non-CpG contexts, the highest 5mC levels were detected in the TTA**CpAC**C and TTC**CpAC**C motifs (**Figure 7B**), consistent with prior findings that DNMT3a, the principal de novo methyltransferase in neurons, preferentially targets CpAC sequences (Lee, Park, and Nakai 2017; Lister et al. 2013); (Mallona et al. 2021). A refined motif logo further revealed a strong preference for T at the −2 position relative to the methylated cytosine (**Figure 7C**). This data also indicates that the modification observed at CpA context is, at least partially, 5mC and not just 5hmC.

**Figure 7:**
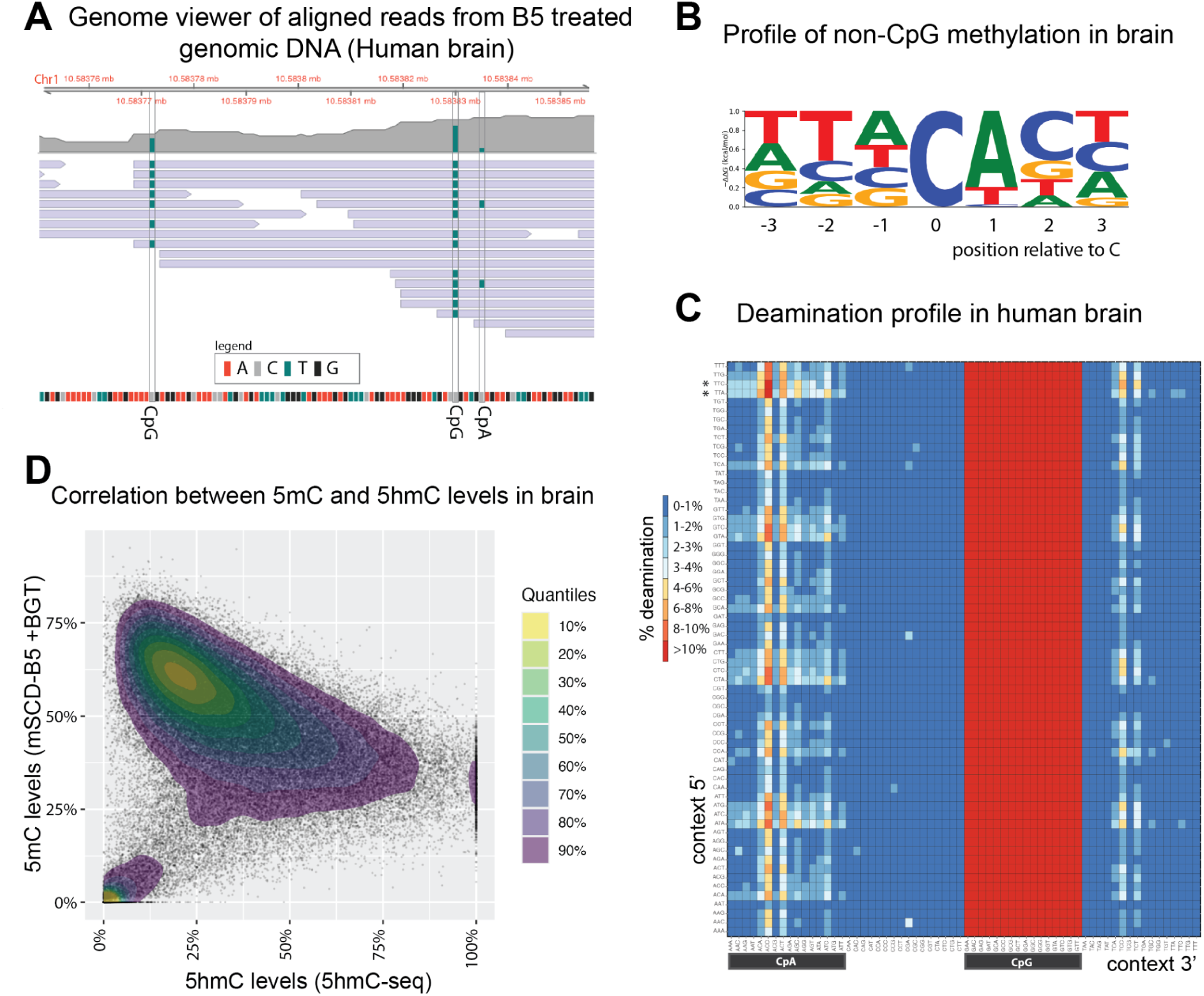
mSCD-B5 deamination analysis reveals high resolution mCpH in human brain genomic DNA. **A)** Genome viewer of aligned reads from mSCD-B5 treated brain genome DNA highlighting base resolution conversion from 5m C-to-T (green) in CpG context and CpA context (human genomic coordinates from GRCh38) **B**) Motif logo of non CpG context preference for methylated sites in the human brain. **C**) Methylation levels measured as the percent deamination at all 5’ contexts (- 3 bases, Y axis) and all 3’ contexts (+3bases, X axis). **D**) Levels of 5mC (y-axis) compared to the level of 5hmC (x-axis) for E. Profile of CpH (left) and CpG (right) methylation.

Another feature of brain genomic DNA is high levels of 5hmC in a CpG context (Globisch et al. 2010). BGT treatment is expected to result in the complete protection of 5hmC revealing 5mC only. We compared the level of 5mC obtained by treatment of mSCD-B5+BGT with the level of 5hmC obtained using NEBNext® Enzymatic 5hmC-seq. The majority of the regions with 5mC have between 15-20% of 5hmC and very few of them have no 5hmC detection at all (**Figure 7D).** This result is in line with the reported amount of 5hmC found in the brain (Wen and Tang 2014). Conversely very few regions without 5mC have detectable 5hmC (**Figure 7D).** Taken together, these results indicate that methylated regions in the brain are almost always a mixture of 5mC and 5hmC.

#### Cell-free DNA methylation

cfDNA fragment length peaks at around 166 base pairs (bp), which corresponds to the combined length of DNA wrapped around a nucleosome (147 bp) plus a 20 bp linker fragment. These fragments result from nuclease degradation and their methylation patterns provide information about the tissue origins (Snyder et al. 2016). Epigenetic marks such as DNA methylation have been used for the detection of fetal DNA (Poon et al. 2002), placenta DNA (Lo et al. 2021; Chim et al. 2005), cancer or organ transplant (K. Sun et al. 2015). Besides methylation, another important epigenetic information contained in cfDNA is nucleosome occupancy. Using enzymatic conversion, recent studies have combined both epigenetic marks in a single experiment (Erger et al. 2020) (Siejka-Zielińska et al. 2021) including TAPS-seq on cf-DNA.

We performed mSCD-B5 treatment together with USER/Q5 on a replicate library from 10ng cfDNA obtained from a healthy donor. We obtained 372 million (for replicate 1) and 304 million (for replicate 2) paired-end reads. The converted library shows the expected fragment distribution sizes (**Figure 8A**). Furthermore, the methylation levels correlate with the fragment sizes (**Figure 8B**) as previously shown. The average deamination on CpG sites is about 60% with a bimodal distribution suggesting full methylated or unmethylated sites (**Supplementary Figure 10A-C**). Replicates have a Pearson’s correlation coefficient of 0.968 (**Supplementary Figure 10D**). In total, we identified methylation on 27.3 million individual CpG sites and 27443 CpG islands, which covers 85.5% total human CpG islands. The methylation profile around CpG island and transcription start sites resembles those of the human NA12878 genomic DNA (**Figure 8C**) (Noë et al. 2024).

**Figure 8:**
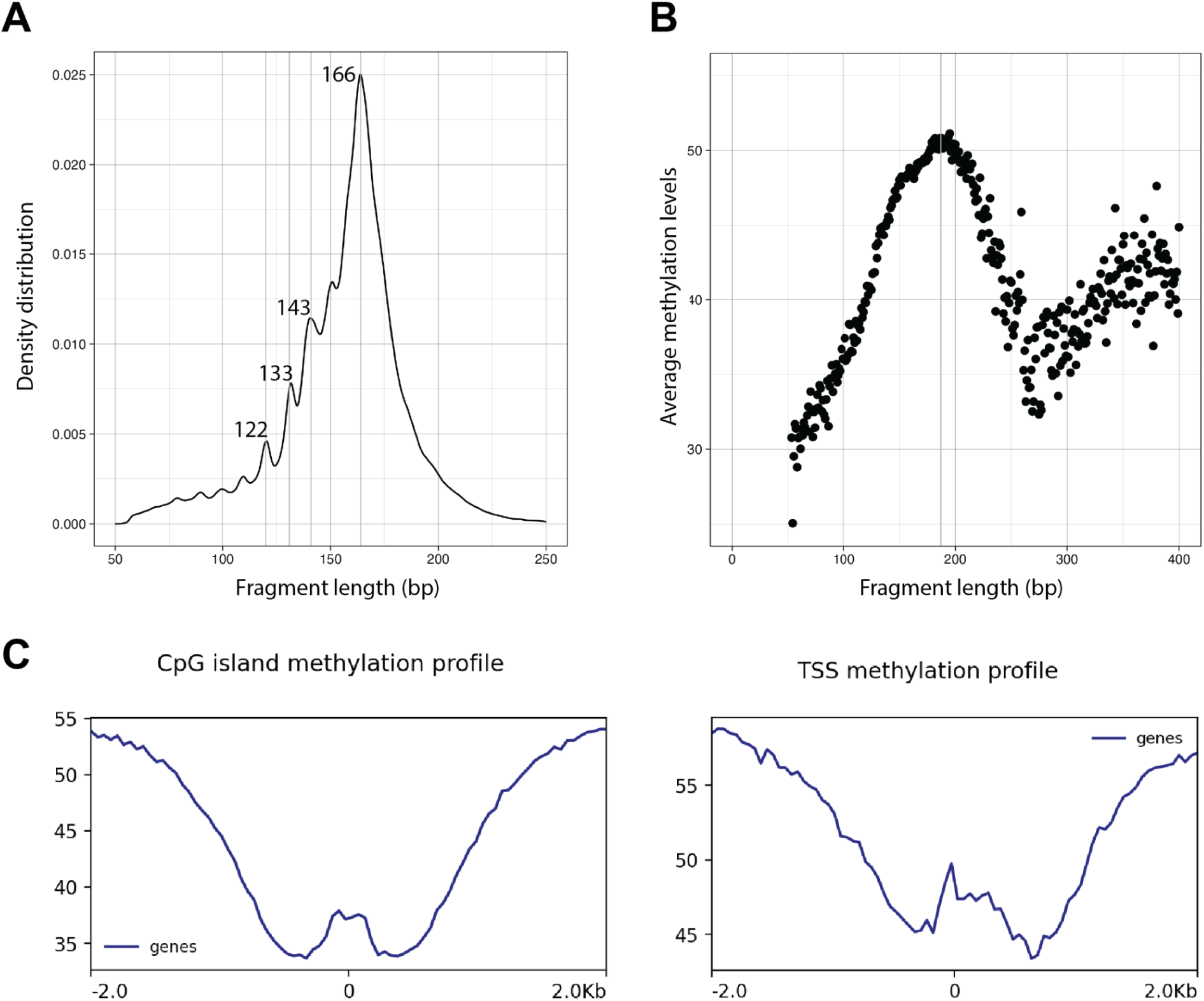
mSCD-B5 sequencing applied to cfDNA. **A)** Fragment size distribution of the converted library. **B)** Average methylation levels function of the insert size (in bp for chr1). The peak methylation level can be found for fragments of ∼187 bp long. **C**) Methylation profile within +/− 2kb region of CpG island (left panel) and Transcriptional Start Site (TSS, right panel).

### 7. Widespread, polyphyletic distribution of mSCD activity in the deoxycytidylate deaminase superfamily

Having characterized mSCD-B5 and its application in methylome sequencing, we sought to address whether mSCD-B5 belongs to a broader functional class of enzymes with similar activities. For this, we searched the IMG/VR virome database (Paez-Espino et al. 2017) for sequences similar to the MetaGPA predicted non-canonical dCMP deaminases. As with MetaGPA, candidate homologs matching our queries were subsequently computationally screened for coassociation with an annotated thymidylate synthase predicted to convert dCMP to either 5m-dCMP or 5hm-dCMP (**Materials and Methods**). The resulting 95 *in silico* identified candidates were added together with the 189 MetaGPA-selected deaminase candidate sequences for further investigation (**Supplementary Figure 9C**).

Candidate deaminases were cloned as synthetic gene constructs into a pET28a *E. coli* expression plasmid with an N-terminal 6x-histidine tag and tested for expression and solubility (see **Materials and Methods**, High Throughput Screening). Of these 284 candidates, 194 could be expressed and detected in the soluble fraction of cell lysates by SDS-PAGE and Coomassie staining. Soluble candidates were bound to magnetic beads functionalized with immobilized nickel, and washed to remove unbound cellular components. Deamination reactions were performed either directly on the beads or after elution by addition of 250 mM imidazole and bead removal with a magnet. The mSCD-B5 deaminase was used as our positive control and baseline for which all the homologs were compared to.

Candidates were screened for methyl specific cytosine deamination on ssDNA *in vitro*, using fluorescently labeled oligonucleotide substrates in a plate-based assay (**Figure 9A-B**). From the 194 soluble candidates, 56 show deaminase activity and 34 have a clear selectivity for 5mC over C. Interestingly, more than 60% of the soluble deaminases from MetaGPA show selective 5mC activity highlighting the strength of our metagenome-wide association study approach in identifying enzymes with specific activities of interest. Next, we performed high throughput sequencing on the gDNA mix libraries treated with the 13 homolog deaminases that demonstrated 5mC selectivity in the CE assay and for which sufficient quantities of purified protein were obtained (**Figure 9A**). In agreement with the CE assay results, the majority of deaminases exhibit higher catalytic activity towards 5mC and 5hmC compared to cytosine (**Figure 9D-E**), with the exceptions of mSCD-GM196 and mSCD-GM197. Two deaminases (MetaGPA2-B4 and MetaGPA2-G10) show minimal or undetectable activity in the NGS assay despite showing activity on CE assay. The highest deaminase activity on 5mC is observed in mSCD-B5 and MetaGPA1-E10, and MetaGPA2-F7 homologs. All active enzymes display varying degrees of activity across different sequence contexts, some with complementary preference compared to B5. For example, MetaGPA2-G12 has preferences for deamination of 5mC in CpG context (**Figure 9D-E**).

**Figure 9:**
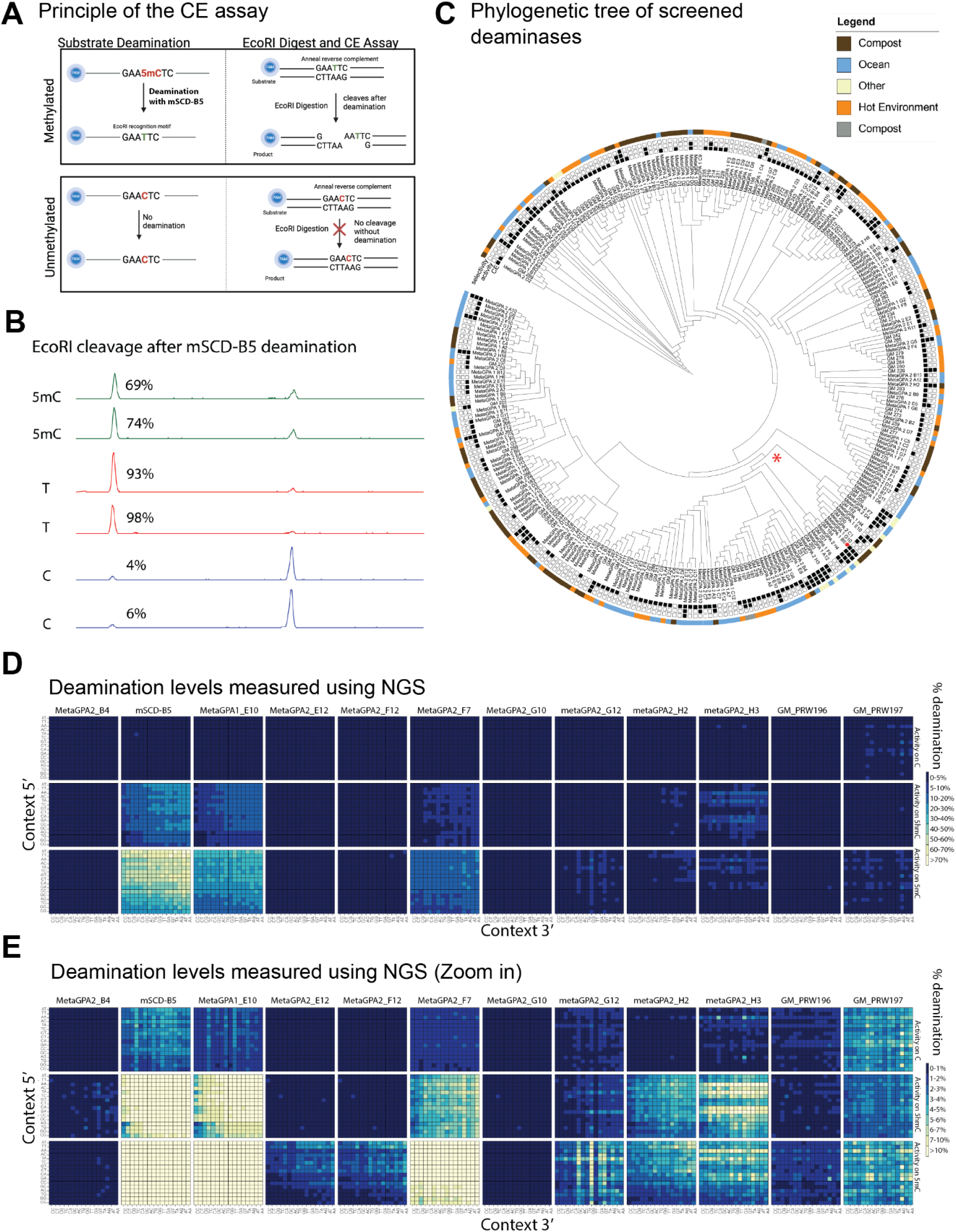
Characterization of homologs of mSCD-B5 on polynucleotides substrate. **A)** principle of the CE assay: The CE Deamination assay uses FAM-labeled oligos containing either GAA5mCTC (L1) or GAACTC (L4) for which deamination of the internal C/5mC creates an EcoRI restriction site. The oligos are annealed to a reverse complement oligo (L3) to form a double-stranded product, followed by digestion with EcoRI. The assay quantifies undigested and digested products using capillary electrophoresis (CE). Control oligos containing GAATTC (L2) or GAAUTC (L5) are processed in parallel to verify the completeness of the EcoRI digestion, ensuring assay reliability. **B)** Deamination activity of mSCD-B5 quantified by CE assay. The two first traces (green) correspond to a CE assay performed in duplicate using the GAA5mCTC (L1) oligonucleotide. The next two traces (red) correspond to a CE assay performed in duplicate using the GAATTC (L2) oligonucleotide (positive control) and the last two traces (blue) correspond to a CE assay performed in duplicate using the GAACTC (L4) oligonucleotide. Deamination levels were estimated using the respective pick areas **C)** Phylogenetic tree of mSCD-B5 homologs generated using Fold Mason Alignment, and built using the UPGMA algorithm with 100 iterations. Boxes are shaded to indicate if the homolog was screened via CE assay, and if there was detectable activity and selectivity on 5mC either through CE or NGS. The outer colored ring denotes the isolation source. The homolog prefixes denote their sources: MetaGPA1 and MetaGPA2 originate from the first and second MetaGPA screens, respectively, while GM represents enzymes identified through genome mining. mSCD-B5 is highlighted on the tree with a red circle. Activity and selectivity on 5mC were quantified by CE Assay. Homologs are colored by isolation sources. **D)** Homolog screen using NGS. Each quadrant corresponds to the deamination levels (in percent from 0 to 100%) at all 5’NNCNN3’ contexts (with N=A, T C or G and C = interrogated cytosine in the reference genome). X and Y axis correspond to contexts 3’ and 5’ to the interrogated C respectively. Percentage deamination on canonical C (Lambda), 5hmC (T4) and 5mC (Xp12) corresponds to top, middle and bottom quadrants respectively. **E)** Same as **D**) with deamination levels (in percent) set from 0 to a maximum of 10% instead.

Taken together, these results demonstrate that the methylation-selective deamination activity on ssDNA is a characteristic of a number of protein sequences annotated as dCMP deaminases. It should be noted that specificity towards deaminating 5mC appears to be a polyphyletic trait. The mixed distribution of 5mC selective deaminases across multiple clades as seen in **Figure 9C** suggests mixed lineages, where no single ancestral event gave rise to a family of sequences having the 5mC specificity. Such a polyphyletic distribution suggests convergent evolution of the trait, with a small number of residues in the active site conferring methylcytosine specificity relative to the canonical activity.

## Discussion

In this study, we use MetaGPA to identify mSCD-B5, a cytosine deaminase with pronounced preference for 5mC and 5hmC over canonical cytosine, in both mononucleotide triphosphates and single-stranded DNA (ssDNA). We find this preference for the 5mC substrate to be sporadically distributed among related members of the dCMP deaminase family (Pfam PF00383) encoded throughout the viral realm. We hypothesize that 5mC-specific deaminases are frequently encoded by bacteriophages which contain 5mC fully replacing C in their DNA and that their biological role is to produce dTTP from deamination of 5mdCTP during infection. This hypothesis is supported by early studies on Xanthomonas phage Xp12 which show that the incorporation of isotopically labeled cytosine into its DNA results in labeling of both 5mC and thymidine (R. Y. Wang and Ehrlich 1982). Xp12 is known to produce 5mdCMP from dCMP by a virally encoded thymidylate synthase homolog expressed during infection. This 5mdCMP is then subsequently phosphorylated to the triphosphate form for uptake by the viral DNA polymerase during genome replication. The deamination of a portion of the 5mdCTP pool produces dTTP and therefore bypasses the need for another thymidylate synthase to convert dUMP to dTMP.

These findings challenge several aspects of the existing deaminase paradigms: first, we demonstrate that, despite being an apparent member of the dCMP deaminase family, mSCD-B5 has a strict requirement for triphosphorylated substrates; Interestingly, the closest known structural homolog to mSCD-B5, the chlorovirus PBCV-1, has activity on both dCMP and dCTP (Zhang et al. 2007). Contrary to PBCV-1, mSCD-B5 has no detectable activity on dCMP; second, we reveal that an enzyme typically specific to mononucleotides can also act on polynucleotide substrates, albeit at lower catalytic rates; and third, we establish that deamination is selectively targeted towards the cytosine derivatives 5mC and 5hmC over the canonical form. Interestingly, mSCD-B5 shows no detectable activity on 4mC, 5fC, caC and glucosylated 5hmC. We have shown here that the lack of activity on glucosylated 5hmC is particularly advantageous for the precise identification of 5mC in genomic DNA containing both 5mC and 5hmC, such as genomic DNA from human brain tissue. In these cases, pre-treatment with BGT protects 5hmC from deamination, enabling specific detection of 5mC.

### For most genomic DNA samples, including those obtained from humans, the majority of bases remain unchanged after conversion

5mC makes up only 1-2% of cytosines in the human genome, and consequently, selective deamination of 5mC mostly keeps the original DNA sequence unchanged. As a result, the converted DNA behaves much like the input DNA, allowing for PCR amplification, target enrichment, and other standard molecular biology techniques that are not feasible with most of the current conversion-based methylome sequencing. Furthermore, sequencing mSCD-B5-treated genomic DNA enhances the ability to accurately map reads to the human genome and facilitates variant calling alongside methylation analysis. This is an important feature of mSCD-B5 specifically for multi-omics applications for which many types of information are layered on top of one another.

### Complete deamination is desirable but not required

Available conversion-based methylome sequencing technologies such as bisulfite sequencing or EM-seq require complete conversion of unmethylated cytosines as residual non-conversion may be erroneously interpreted as methylation. Under certain conditions, such as prolonged incubation times with converting reagents, these same methods can also “over-convert” and deaminate 5mC, resulting in loss of signal. In contrast, the direct deamination of 5mC by mSCD-B5 or enzymes with similar properties does not require complete conversion since the background unmethylated cytosine does not interfere with the readout. Furthermore, given a demonstrated linear correlation between methylation levels and 5mC conversion, methylation levels can be accurately modeled even with incomplete conversion. However incomplete conversion by mSCD-B5, which has been typically found to be around 20% may still produce false negatives depending on a combination of factors such as sequencing depth and absolute methylation levels.

### Deamination of 5mC produces T, a canonical nucleobase in DNA

mSCD-B5 deamination of 5mC results in the canonical base T, which is compatible with standard molecular biology protocols and subsequent enzymatic treatments that may be hindered or blocked by uracil, such as certain DNA or RNA polymerases, restriction enzymes, and genome editing enzymes. Amplification is therefore not necessary as mSCD-B5 treated libraries should be fully compatible with direct PCR-free sequencing. Although mSCD-B5 has a background C to U conversion of around 1-4% in our hands, products of this undesired activity can be eliminated by treating converted libraries with USER while leaving intact the product of deamination of 5mC. However, under these conditions, the amount of starting material for amplification and fragment sizes is inversely correlated with the background deamination rate on C that cannot exceed 5% using standard library preparation. Single-stranded ligation of adapters after conversion, may be an alternative strategy to avoid loss of material.

We have demonstrated that USER treatment enhances the sensitivity of methylation detection down to 0.1%. This level of sensitivity not only reveals previously undetected methylation in standard controls used in methylome sequencing but also enables the detection of low-abundance methylation events either difficult to achieve or beyond the reach of existing methods. This sensitivity holds significant potential for applications such as clinical diagnostics, where ultra-sensitive methylation detection is essential for early disease biomarker identification.

## Materials and Methods

### Genomic DNA

The *E. coli* K-12 DHB4, phage Xp12, phage T4gt genomic DNA used in this study were obtained from New England Biolabs (NEB). The mCpG plasmid pUC19 was obtained from NEB. The unmethylated lambda phage genomic DNA was purchased from Promega (catalog number: D1521). Human GM12878 genomic DNA was purchased from Coriell Institute (catalog number: NA12878). Human brain genomic DNA from a single donor was purchased from biochain (catalog number: D1234035, LOT: C607510). For cfDNA, the single donor human blood derived plasma was purchased from Innovative Research (catalog number: IPLASK2E2ML). DNA extraction for cfDNA was performed using biochain cfPurer-V22-cfDNA extraction kit (catalog number: K5011625-V2). We purchased HCT116 WT cells from ATCC (catalog number: CCL-247). The DNMT1 and DNMT3b DKO HCT116 strain was a gift from Bert Vogelstein (John Hopkins Medicine). Genomic DNA extraction from HCT116 WT and DKO cells were performed with Monarch genomic DNA purification kit (NEB, catalog number: T3010), following the protocol for genomic DNA extraction from cells. The *dcm-6* (ER3662, CGSC Strain#: 6475, fhuA2 or fhuA31, lacY1 or lacZ4, tsx-1 or tsx-78, glnX44(AS), galK2(Oc), galT22, dcm-6, dam-3, mtlA2, metB1, thiE1), *Δdcm* (ER2861, fhuA2::IS2 Δ(lacZ)4826 glnX44 e14-trpE31 Δ(hisG)1 rpsL104(StrR) xyl-7 mtlA2(Fs) metB1(FS) Δ([fimB or yjiT]-opgB)114::IS10 Δ(dcm::FRT1)1) and the *dcm+* (DHB4, GenBank accession: CP014270.1) genomic DNA were obtained from NEB. The m4C containing genomic DNA from *Natrinema pallidum* BOL6-1 archaea was also obtained from NEB.

### MetaGPA screen

We performed two MetaGPA screens in this study. The first MetaGPA screen method using TET/APOBEC/USER was described previously (Yang et al. 2021). For the same purpose of enriching cytosine modified phages in the microbiome, we conducted a separate screen using treatment with a restriction enzyme (RE) cocktails containing Hpa II, Hae III, Alu I, Aci I, Bfa I, Hha I, HpyCH4IV, HpyCH4V and Nla III. All enzymes are products of NEB and have been tested to be sensitive to 5mC (Flodman et al. 2020) and therefore can be used to digest phage genomes with canonical cytosine. First, microbiome phage fraction genomic DNA described in (Yang et al. 2021) was split into two for RE cocktail treatment or untreated control. For the treatment group, 1 µl of each enzyme cocktail component was added to the DNA sample and the digestion reaction was incubated at 37°C for 30 min in 1x rCutSmart Buffer (NEB, catalog number: E6004). Water was added to the untreated control for the digestion reaction. The digestion reaction was stopped by adding 5µl Proteinase K (NEB, catalog number: P8107) and incubation at 37°C for 30 min, followed by column purification using Zymo oligo clean and concentrator kit (catalog number: D4060). Second, DNA samples were incubated with AMPure XP beads (Beckman Coulter, A63880, dilute to 35% v/v) at a beads to sample ratio of 3.5:1 to retrieve large fragments (greater than 3kb). The resulting DNA fragments were eluted in 50 µl elution buffer and sheared to 250bp using Covaris S2 Focused Ultrasonicator. Preparation of DNA libraries from above samples was done following instructions of NEBNext Ultra II DNA library preparation kit (catalog number: E7645). Then, a second round of RE cocktail treatment was conducted with DNA libraries in reaction condition as described above. After DNA purification using Zymo oligo clean and concentrator kit, final products were indexed (with NEBNext Multiplex Oligos for Illumina, catalog number: E6440), amplified using NEBNext Ultra II Q5 Master Mix (catalog number: E7645) and pooled for sequencing on an Illumina NextSeq instrument with paired end reads of 75bp.

### Deaminase *in vivo* screen

We cloned the seven deaminase candidates into pET28a backbone with N term Histag. The plasmids bearing the deaminase genes were used to transform T7 Express Competent E. Coli (#C2566, NEB) together with the 5hmC or 5mC incorporation plasmids (5hmdCpre or 5mdCpre) to examine the *in vivo* deaminase activities. For each transformed strain, a single colony transformant was used to inoculate 0.5 mL terrific medium in a well of a 96-well deep well plate with kanamycin and chloramphenicol antibiotics at the concentrations of 50 and 34 µg/mL, respectively. The 96-well plate was incubated at 37°C with 800 rpm shaking until OD_600_ reached 0.4-0.6, the plate was cooled to RT, then induced with 0.1 µM IPTG and further incubated with shaking at 18 °C for 15 hours. The cultures were pelleted and subjected to plasmid DNA isolation using Monarch Plasmid Miniprep Kit (NEB) following the manufacturer’s protocol. The isolated plasmid DNA was hydrolyzed to nucleosides using the Nucleoside Digestion Mix (NEB) performed at 37°C for >2 hours. The nucleoside solutions were then filtered through a hydrophilic PTFE 0.2 µm centrifugal filter, where the flowthroughs were subjected to HPLC-MS analysis for the nucleoside composition. Methods described at the LC-MS section were used for the analysis.

### Protein Expression and Purification

About 50 ng of plasmid DNA was transformed into T7 Express lysY/Iq Competent *E. coli* (#C3013, New England Biolabs, Ipswich, MA, USA), plated onto LB-Kan agar and incubated at 37°C overnight. Single colonies were inoculated into LB with kanamycin (40 µg/mL). The culture was incubated at 37°C, with shaking at 225 rpm until OD_600_ reached 0.4-0.6. Cells were then induced with IPTG (final concentration 0.5 mM) and incubated overnight at 22°C with 225 rpm shaking. The following day cells were harvested and the expressed protein was purified using IMAC resin. Enzymes were then stored at −20°C until further use. Experiments monitoring mononucleotide deamination activity by LC-MS were done with proteins from a similar purification procedure except that the one column procedure using HisTrap was replaced by two columns procedure using His-Trap followed by HisTrap Heparin.

### Homolog deaminase candidate selection

A phylogenetic tree of homologs selected for screening was constructed to visualize relationships between sequence similarity and empirically tested activity. Protein sequences for all homologs were run through AlphaFold2 to predict protein structures with 12 recycles and 5 model outputs. Best scoring models for each sequence were used to guide protein alignments using FoldMason under default settings. A phylogenetic tree was generated using an unweighted pair group method with arithmetic mean (UPGMA) algorithm via PHYML with 100 iterations.

### Homolog deaminase plasmid cloning

Homologs selected from the study were cloned into pET28a backbone with N term Histag. The recombinant plasmids were ordered through Twist Bioscience.

### High Throughput expression and screening of homolog deaminases

For rapid, high-throughput screening of mSCD-B5 homologs, a semi-automated 96-well plate approach was performed. Plasmid constructs were purchased from GenScript Biotech or Twist Bioscience using the pET28a plasmid backbone (Piscataway, NJ, USA). NEB® C3010 T7 Express lysY Competent *E. coli* (High Efficiency) or NEB® C3013 T7 Express lysY/Iq Competent *E. coli* (High Efficiency) (New England Biolabs, Ipswich, MA, USA) were transformed with ∼5 ng of plasmid using the 5-minute transformation protocol, according to the manufacturer’s instructions. Transformation reactions were then transferred to deepwell 96-well plates containing room temperature NEB SOC Outgrowth Media (New England Biolabs, Ipswich, MA, USA), and incubated for 1 hour at 37°C with shaking at 800 rpm. Cells were then subcultured into 2 mL deepwell 96-well plates containing 1 mL Dynamite media (Taylor, Denson, and Esposito 2017) with kanamycin (40 µg/mL) per well. Subcultured cells were returned to 37°C for 2-4 hours with shaking at 800 rpm. Cells were then induced with 0.5 mM IPTG and incubated 16-18 hours at 22°C with shaking at 800 rpm.

The following day proteins were purified using a KingFisher Flex (ThermoFisher Scientific, Waltham, MA, USA). Cell pellets were resuspended in IMAC binding buffer (50 mM sodium phosphate, pH 8.0, 300 mM sodium chloride, 10 mM imidazole) for 25 minutes at 25°C, shaking at 1200 rpm. To lyse cells, BugBuster Master Mix (Millipore, Burlington, MA, USA) or NEBExpress® *E. coli* Lysis Reagent (New England Biolabs, Ipswich, MA, USA) was added and incubated for 30 minutes at 25°C, 1200 rpm. Proteins were captured using 50 µL of NEBExpress® Ni-NTA Magnetic Beads per well and washed using the typical reaction protocol described by the manufacturer. Proteins captured on nickel beads were ultimately resuspended in activity buffer (50 mM bis-tris, pH 6.5) and immobilized proteins were screened for activity using the CE assay (described above).

### Thymidylate synthase expression and substrate activity confirmation *in vivo* assay

Each cognate thymidylate synthase gene of the seven mSCD hits was synthesized from Genscript and was cloned to a pET28b(+) vector under the control of a T7 promoter. Each thymidylate synthase construct was used to transform the T7 express E coli strain harboring the nucleotide monophosphate kinase from bacteriophage T4 cloned in a pACYC vector. An in vivo assay was performed using the transformed strains to examine the thymidylate synthase activity. Starting cultures were prepared from single colony inoculation of the transformed strain and were cultured at 37°C to stationary phase. The starting cultures were used to inoculate the LB media with kanamycin (50 µg/mL) and chloramphenicol (25 µg/mL) antibiotics at 1 in 20 dilutions. The cell cultures were then incubated at 37°C with agitation until an OD600 reached the mid-log phase before adding the isopropyl beta-D-1-thiogalactopyronoside (IPTG) to a 100 µM final concentration. The cultures were then returned to the incubation at 37°C until stationary phase. Next, we pelleted the culture cells and isolated the plasmid from the culture to examine the nucleotide composition. A Monarch plasmid miniprep kit (NEB, Cat# T1110) was used to isolate the plasmids following the manufacturer protocol in which the afforded plasmids were further treated with the Nucleotide digestion mix (NEB, Cat# M0649) for hydrolysis to nucleosides. The nucleoside mixtures were then passed through the 0.2 µm PTEF centrifuge filter, and the filtrates were subjected to the HPLC-MS analysis as the methods described in the HPLC-MS section.

### CE-assay

We devised a capillary electrophoresis (CE) assay to test for activity on single stranded DNA (CE assay). Single-stranded 31 nucleotide DNA oligos were designed and ordered from IDT with a 5’FAM label and an EcoRI (GAAXTC) restriction site at 12^th^ – 17^th^ position, where X was replaced with a 5mdC (L1), dT (L2), dC (L4) or dU (L5). Single stranded oligos were incubated with mSCD-B5 at a 30:1 enzyme:substrate ratio in 50mM Bis-Tris, pH 6, 100uM ZnCl_2_ at 42°C for 1 hour. After deamination, the unlabeled reverse complement to the sequence was annealed to the oligos at 95°C for 10 min, then slowly cooled to room temperature. Reactions were cleaned with Zymo Oligo clean and concentrate columns, and eluted, then were digested with EcoRI-HF (NEB #R3101) for 1hr at 37°C in rCustmart® buffer (NEB #B6004S) following manufacturer recommendations. Reactions were stopped by the addition of 1µl of ProteinaseK (NEB #P8107S) and incubated at room temperature for 15 minutes. Finally oligos were diluted to 2nM and submitted for Capillary Electrophoresis on Applied Biosystems 3730xl DNA Analyzer machine running POP6 polymer. Peaks were analyzed with Ebase (NEB).

Oligo sequence are listed below:

L1: /56-FAM/-ATACACCGATGCAGAA**/iMe-dC/**TCATGGGCTTACGC,
L2:/56-FAM/-ATACACCGATGCAGAA**T**TCATGGGCTTACGC,
L3:GCGTAAGCCCATGAATTCTGCATCGGTGTAT,
L4:/56-FAM/-ATACACCGATGCAGAA**C**TCATGGGCTTACGC,
L5:/56-FAM/-ATACACCGATGCAGAA**/ideoxyU/**TCATGGGCTTACGC

### General instrumentation of liquid chromatography (LC) and liquid chromatography-mass spectroscopy (LC-MS)

LC-MS was performed on an Agilent 1290 Infinity II UHPLC-MS system equipped with a G7117 Diode Array Detector and an LC-MSD XT G6135 Single Quadrupole Mass Detector. Chromatography performed with the instrument was on a Waters XSelect HSS T3 C18 column (4.6 × 150 mm, 3 µm particle size) and operated at a flow rate of 0.5 mL/min with a binary gradient mobile phase consisting of 10 mM ammonium acetate (pH 4.5) and methanol. The course of chromatography was monitored at 260 nm. Mass spectrometry was operated in both positive (+ESI) and negative (-ESI) electrospray ionization modes. MS was performed with a capillary voltage of 2500 V at both modes, a fragmentor voltage of 70 V, and a mass range of m/z 100 to 1000. Agilent ChemStation software was used for the primary LC-MS data processing. ChemStation produced chromatograms were further annotated in Adobe Illustrator.

### Deaminase assay on mono-nucleotides,single-strand DNA oligo

In the deamination reaction on single-strand DNA oligo, 3 µM synthesized ssDNA oligo with an internal 5mC, 5hmC, 5fC and 5caC were incubated with 5.8 µM deaminase, 0.5 mg/mL rAlbumin (NEB, catalog number: E9200), 0.1 mM ZnCl_2_ and 1x Deamination Reaction Buffer (#E3356, NEB) at 42°C for 1 hour. Final products were purified with Zymo oligo clean and concentrator kit, resuspended in 17 µl water and proceeded with nucleoside digestion reaction (NEB, catalog number: M0649) to prepare for LC-MS.

5gmC single strand oligo was not available through commercial oligo synthesis services. To prepare it, we first annealed an equal molar of 5hmC top strand (5hmC ss oligo) and bottom strand by heating the mix to 95°C for 5 min and gradually cooling it down to room temperature. The resulting double stranded oligos were then incubated overnight with T4 Phage β-glucosyltransferase (BGT) at 37°C following the manufacturer’s instruction (NEB, catalog number: M0357S) to convert 5hmC on top strand to 5gmC. We used Norgen Biotek’s oligo cleanup and purification kit (SKU 34100) to clean up the 5gmC double strand oligo. Finally, 5gmC single strand oligos were prepared by heat denaturation at 95°C for 10 min and immediately moving to an ice bath for 3 min.

Oligo sequence are listed below:

5mC ss oligo: 5’-ACACCCATCACATTTACAG{**5mC**}CGGAAAGAGTTGAATGTAGAGTTGG-3’
5hmC ss oligo: 5’-ACACCCATCACATTTACAG{**5hmC**}CGGAAAGAGTTGAATGTAGAGTTGG-3’
5fC ss oligo: 5’-ACACCCATCACATTTACAG{**5fC**}CGGAAAGAGTTGAATGTAGAGTTGG-3’
5caC ss oligo: 5’-ACACCCATCACATTTACAG{**5caC**}CGGAAAGAGTTGAATGTAGAGTTGG-3’
Bottom strand: 5’-CCAACTCTACATTCAACTCTTTCCGGCTGTAAATGTGATGGGTGT-3’

### NGS library preparation with mSCD-B5 deamination

To prepare the library for sequencing, *E. coli*, Human GM12878 or human brain genomic DNA (see Genomic DNA section) was sheared to 300 bp in 130 μL of TE buffer (10 mM Tris pH 7.5, 1 mM EDTA) using Covaris S2 Focused Ultrasonicator. Sheared spike-in phage control genomic DNA (containing Lambda phage, Xp12 and T4gt) and/or mCpG pUC19 plasmid DNA were added when desired. One reaction of NEBNext Ultra II DNA Library Prep Kit for Illumina (NEB #E7645) was used for each sample. The sample was End-prepared and ligated to regular adaptor following the manual (NEB #E7645). The DNA library was purified with 1x volume of NEBNext Sample Purification Beads (NEB #E7103) and eluted with 16 μL of water.

To denature the dsDNA, 16 μL prepared library sample was mixed with 4 μL freshly prepared 0.1N NaOH and incubated in a preheated thermocycler at 85°C for 10 minutes. After the heat denaturation, the reaction was placed immediately on ice for 3min. Deamination reaction was conducted in 50 μL volume with 20 μL denatured DNA library, 5.8 µM deaminase, 0.5 mg/mL rAlbumin (NEB, catalog number: E9200), 0.1 mM ZnCl_2_ and 1x Deamination Reaction Buffer (#E3356, NEB) at 42°C for 1 or 2 hours. The deamination reaction was inactivated by incubating with 2μL proteinase K at 37°C for 30 minutes. The reaction was purified with 1x volume of NEBNext Sample Purification Beads (NEB #E7103) and eluted with 40 μL of 0.1xTE buffer. Libraries were indexed with NEBNext Multiple Oligos for Illumina (NEB #E6440) and amplified using either NEBNext Ultra II Q5U Master Mix (NEB #M0597) or NEBNext Ultra II Q5 Master Mix (NEB #M0544).

### Methylation sensitivity experiment

A sample containing 4% methylation at dcm sites was obtained by mixing dcm+ genomic DNA (100% methylation at dcm sites) with Δdcm genomic DNA (0% methylation) at a 4:96 molar ratio. Serial 1:2 dilutions were performed by mixing an equimolar amount of Δdcm genomic DNA to the previous dilution, starting from the 4% sample. The serial dilutions generated samples containing methylation levels at 2%, 1%, 0.5%, 0.25%, 0.13%, 0.06% and 0.03% respectively. Each dilution level sample was split into two duplicates.

The methylation levels were independently quantified for the 100% and 4% dcm+ sample using LC-MS/MS. The 5mC levels were determined to be 1.2% and 0.03% of total C for the 100% and 4% dcm+ sample respectively. This quantification is consistent with the fact that only ∼1% of the C are methylated in the E.coli genome and is in line with the targeted 100% and 4% methylation at C<u>C</u>WGG sites.

### NGS library preparation with mSCD deamination on cfDNA

10 ng of extracted cfDNA was directly mixed with sheared spike-in control genomic DNA. The library preparation was performed using one reaction of NEBNext Ultra II DNA Library Prep Kit for Illumina (NEB #E7645). We used the diluted adaptor (1.5 μM) for the ligation step and followed the manual for the remaining steps of library preparation. The prepared DNA library was processed the same as previously described for deamination reaction and PCR amplification.

Preparation of synthetic oligos and *E.coli* samples for residual methylation detection

For the methylated oligos experiment, we synthesized oligos containing one or no internal 5mC site from IDT (Integrated DNA Technologies). We also designed adaptor sequences on both ends of the oligo to facilitate illuminate sequencing. The forward and reverse oligos were annealed to make dsDNA by mixing equal molarity of each, heating to 95°C for 4 minutes and gradually cooling on the benchtop for about 20 minutes. We then used the fully methylated oligo (100% methylation, methylated F + methylated R) and the fully unmethylated oligo (0% methylation, unmethylated F + unmethylated R) to prepare samples with serial dilution to obtain methylation levels at 80%, 40%, 20%, 10%, 5%, 2%, 1% and 0.5%. The deamination reaction on each sample was conducted as previously described. PCR amplification was performed with NEBNext Ultra II Q5 Master Mix (NEB #M0544) and NEBNext Multiple Oligos for Illumina (NEB #E6440). The sequence of oligos are listed below:

Unmethylated F: 5’CACTCTTTCCCTACACGACGCTCTTCCGATCTGGACACAGGGGTTGCCCAGGCTACAGATCCGGGTTCAGGCCCAGCCGTGGGGACTGTTCACCTCTGAGAGGGCAGCCCTGCCCCGCGGCCCTGGCCCTGCCCTGCACCCGGCTGGGTCTCATCTGACACGCAGATCGGAAGAGCACACGTCTGAACTCCAGTCAC
Unmethylated R: 5’GTGACTGGAGTTCAGACGTGTGCTCTTCCGATCTGCGTGTCAGATGAGACCCAGCCGGGTGCAGGGCAGGGCCAGGGCCGCGGGGCAGGGCTGCCCTCTCAGAGGTGAACAGTCCCCACGGCTGGGCCTGAACCCGGATCTGTAGCCTGGGCAACCCCTGTGTCCAGATCGGAAGAGCGTCGTGTAGGGAAAGAGTG
Methylated F: 5’CACTCTTTCCCTACACGACGCTCTTCCGATCTGGACACAGGGGTTGCCCAGGCTACAGATC/**iMe-dC**/GGGT TCAGGCCCAGCCGTGGGGACTGTTCACCTCTGAGAGGGCAGCCCTGCCCCGCGGCCCTGGCCCTGCCCTGCACCCGGCTGGGTCTCATCTGACACGCAGATCGGAAGAGCACACGTCTGAACTCCAGTCAC
Methylated R: 5’GTGACTGGAGTTCAGACGTGTGCTCTTCCGATCTGCGTGTCAGATGAGACCCAGCCGGGTGCAGGGCAGGGCCAGGGCCGCGGGGCAGGGCTGCCCTCTCAGAGGTGAACAGTCCCCACGGCTGGGCCTGAACC/**iMe-dC**/GGATCTGTAGCCTGGGCAACCCCTGTGTCCAGATCGGAAGAGCGTCGTGTAGGGAAAGAGTG

For *E. coli* experiment, we first prepared a 4% methylation sample by mixing 4% of the sheared *dcm+* DHB4 strain with 96% of the sheared *Δdcm* ER2861 strain. Then, a 2 fold serial dilution was made by mixing the methylated sample with*Δdcm* ER2861 strain to get 2%, 1%, 0.5%, 0.25%, 0.13%, 0.06% and 0.03% methylation samples. Samples with different methylation levels were prepared in duplicates. Each sample were subjected to one reaction of NEBNext Ultra II DNA Library Prep Kit for Illumina (NEB #E7645). Deamination reactions were conducted as described before. PCR amplification was conducted with NEBNext Ultra II Q5 Master Mix (NEB #M0544) and USER treatment.

### BGT treatment

To obtain 5mC specific deamination, prior to deamination reaction, the prepared DNA library (16 μL) was treated with 1x CutSmart buffer, 0.4 μL UDP-Glucose (2mM), 1 μL T4-BGT (10U/μL) in a 50 μL reaction (NEB #0357) and incubated at 37°C for 1 hour. The reaction was purified with 1.8x volume NEBNext Sample Purification Beads (NEB #E7103) and eluted in 16 μL water for downstream deamination reaction.

### USER treatment

2 μL USER enzyme (NEB #M5505) was added to each PCR amplification reaction containing deaminated DNA library, NEBNext Multiple Oligos for Illumina (NEB #E6440) and NEBNext Ultra II Q5 Master Mix (NEB #M0544). An additional 15 min incubation at 37°C was performed before the PCR reaction.

### Tet-assisted Pacbio sequencing

1 μg of *E. coli* DNA was sheared to an average size of 5-10KB using gTubes (Covaris, Woburn, MA) and concentrated using 0.6V Ampure beads (Pacific Biosciences). Libraries were prepared using SMRT-bell Express Template Prep kit 2.0 (PacBio, #100-938-900) according to the manufacturer’s protocol. Barcoded libraries were TET-treated following an abbreviated protocol for NEBNext Enzymatic Methyl-Seq Conversion Module (NEB, #E7120) to enzymatically convert modified cytosines into 5-carboxylcytosine. Briefly, libraries were incubated for 1 hour at 37°C with TET2 in the presence of TET2 reaction buffer, an oxidative supplement, DTT, and an oxidation enhancer followed by addition of stop reagent and incubation at 37°C for 30 minutes. TET2 treated libraries were cleaned up using 1V Ampure beads (Pacific Biosciences).

TET-treated, barcoded libraries were prepared for sequencing on PacBio Sequel II using Sequel II sequencing kit 2.0 (PaBio, #101-820-200). Pacific Biosciences SMRTlink web portal was used to calculate sequencing conditions and perform post-sequencing analysis.

### Bioinformatics analysis

1. Substitution analysis (Supplementary Figure damage). We used a similar approach as described in (Baum et al. 2021). Briefly, mapped reads were partitioned based on their origin (either Read 1 or Read 2 from paired-end reads) using samtools (version 1.8) using flagstat -f 64 (for Read 1) and -f 128 (for Read 2). Samtools mpileup (version 1.8) was subsequently applied to the split read files with the following parameters: --a -O -s -q 10 -Q 30. Reverse reads were reoriented in the forward direction, and the fractions of all 12 substitutions (C>A, C>T, C>G, G>A, G>C, G>T, A>T, A>G, A>C, T>A, T>C, T>G) were computed for all paired-end read1 across all possible dinucleotide contexts.
2. Sensitivity of mSCD-B5 in NGS assay. Reads were mapped to the composite genome containing the DHB4 E.coli genome (NZ_CP014270.1). Variants relative to the *E. coli* DHB4 genome were identified using Snippy (snippy 4.4.0), based on mapped reads from the unmethylated Δdcm strain and removed from further analysis (780 genomic positions in total) to avoid interference with C>T substitution from deamination. The fraction of C>T substitution for both paired-end read1 and 2 across all possible NCNNN contexts were obtained, as described in the Substitution Analysis section. To identify imbalances indicative of deamination (Baum et al. 2021; Chen et al. 2017), the fraction of substitutions in Read 2 was subtracted from that in Read 1.
3. Free circulating DNA. To obtain methylation on a read level, we used Biscuit (Zhou et al. 2024) with the following parameters : biscuit cinread -p “QNAME, QPAIR, STRAND, BSSTRAND, MAPQ, QBEG, QEND, CHRM, CRPOS, CGRPOS, CQPOS, CRBASE, CCTXT, CQBASE, CRETENTION” -g chr1 -t cg on the mapped reads. Methylation profile around +/−2kb CpG islands and transcription start sites were obtained with deeptools: computeMatrix--referencePoint center -a 2000 -b 2000 and plotProfile
4. *In silico* identification of candidate MSCDs Candidate mSCD-B5 homologs were identified for screening in four steps. The first two steps were performed using the Domainator, an open-source software suite which contains several tools for comparative genomics. First, the “Earth’s metavirome” database, IMG-VR (Camargo et al. 2023), was searched for contigs containing matches to the Pfam profile “Cytidine and deoxycytidylate deaminase zinc-binding region”(PF00383). Contigs containing matches above eval: 10^-7^ and greater than 3 kb in length were recovered. Second, contigs which lacked a match to “Thymidylat_synth” (PF00303) at eval greater than 10^-7^ were discarded. Third thymidylate synthase homologs, together with the ThyA sequence of *E. coli* MG1655 were aligned using Muscle. Only contigs containing a thymidylate synthase homolog encoding an aspartate at positions homologous to N177 in the *E. coli* sequence were retained. Lastly, 234 (check this number) unique cytidine deaminases from these contigs were aligned together with “landmark” cytidine deaminases from a variety of cellular and viral sources and the resulting MSAs were used to generate phylogenetic trees with PhyML (Guindon et al. 2010). Ninety-five putative methyl-specific cytidine deaminases were arbitrarily selected from each of the resulting clades obtained in the dendrogram.. This search resulted in 95 candidate deaminases.
5. Structural similarity search : A search for proteins with structures similar to mSCD-B5 was done first predicting the mSCD-B5 structure using AlphaFold2 (Jumper et al. 2021) and querying the resulting best model against the PDB using Foldseek (van Kempen et al. 2024), which returned the following matches: 7fh4_B; e-value: 2.92 e-15 followed by PBDs 4p9e, 4p9d, 4p9c, 1vq2 with e-values: 1.65 e-9, 1.88 e-9, 1.65 e-9, 2.28 e-9, respectively.
6. MetaGPA analysis: MetaGPA code and analysis is available as described in (Yang et al. 2021). For this study, we used default parameters in MetaGPA main pipeline except for adjusting the contig enrichment score to 10 (-C 10). The Pfam database release 33.0 was used for annotation analysis.
7. Phylogenetic tree analysis with annotated dCMP deaminase from MetaGPA: Full coding sequences of annotated dCMP deaminases from both MetaGPA identified modified and unmodified contigs were extracted with custom python script. Multiple sequence alignments were performed with mafft v7.508 (Katoh and Standley 2013). The resulting aligned fasta files were subjected to construct phylogenetic trees using the maximum likelihood method in the phylogenetic analysis program RAxML v8.2.12 (Stamatakis 2014). We chose the -f a option to do rapid bootstrap analysis and the -m PROTGAMMAAUTO model to automatically determine the best protein substitution model to be used for the dataset. The parsimony trees were built with random seeds -p 1237 and -x 1234. The online tool iTOL (https://itol.embl.de/) was used to visualize trees.
8. Mapping tools and methylation detection analysis: If not specified otherwise, we used BWA-MEM (H. Li and Durbin 2010) for mapping of reads to bacterial or human genomes. Reads mapping to Xp12 and T4gt genomes was performed with bwameth.py (arxiv.org/abs/1401.1129). Methylation on single base resolution was analyzed and detected with Methyldackel (https://github.com/dpryan79/MethylDackel) or custom script.
9. Pacbio sequencing data processing and analysis: Pacbio long reads were processed, demultiplexed and ccs reads were generated with smrtlink portal. We then performed base modification analysis to obtain per base modification IPD. Data was analyzed and plotted with custom python script.

## Acknowledgments

We would like to thank the NEB Sequencing core and IT for providing support. Chen Song for cfDNA, Katherine O’Toole for modified Cytosine oligos, Brian Anton and Julie Beaulieu for dcm +/− E. coli strains, Pierre Esteve and Karthikeyan Raman for HCT116 cells. Louise Williams and Daniel Evanich for Brain genomic DNA. Brad Langhorst, Lana Saleh, Sriharsa Pradhan for suggestions. Sean Lund for cloning suggestions. Andy for protein purification. Jodi Cook and Brett Gent for reagent ordering. Shuang-Yong Xu for advice on the restriction enzyme cocktail, Alexey Fomenkov for advice on the Pacbio experiment setup. Tom Evans and Nicole Nichols for manuscript suggestions.

## Data and code availability

All raw and processed sequencing data generated in this study have been submitted to the NCBI Gene Expression Omnibus (GEO; https://www.ncbi.nlm.nih.gov/geo/) under accession number.

## Figures

**Supplementary Figure 1:**
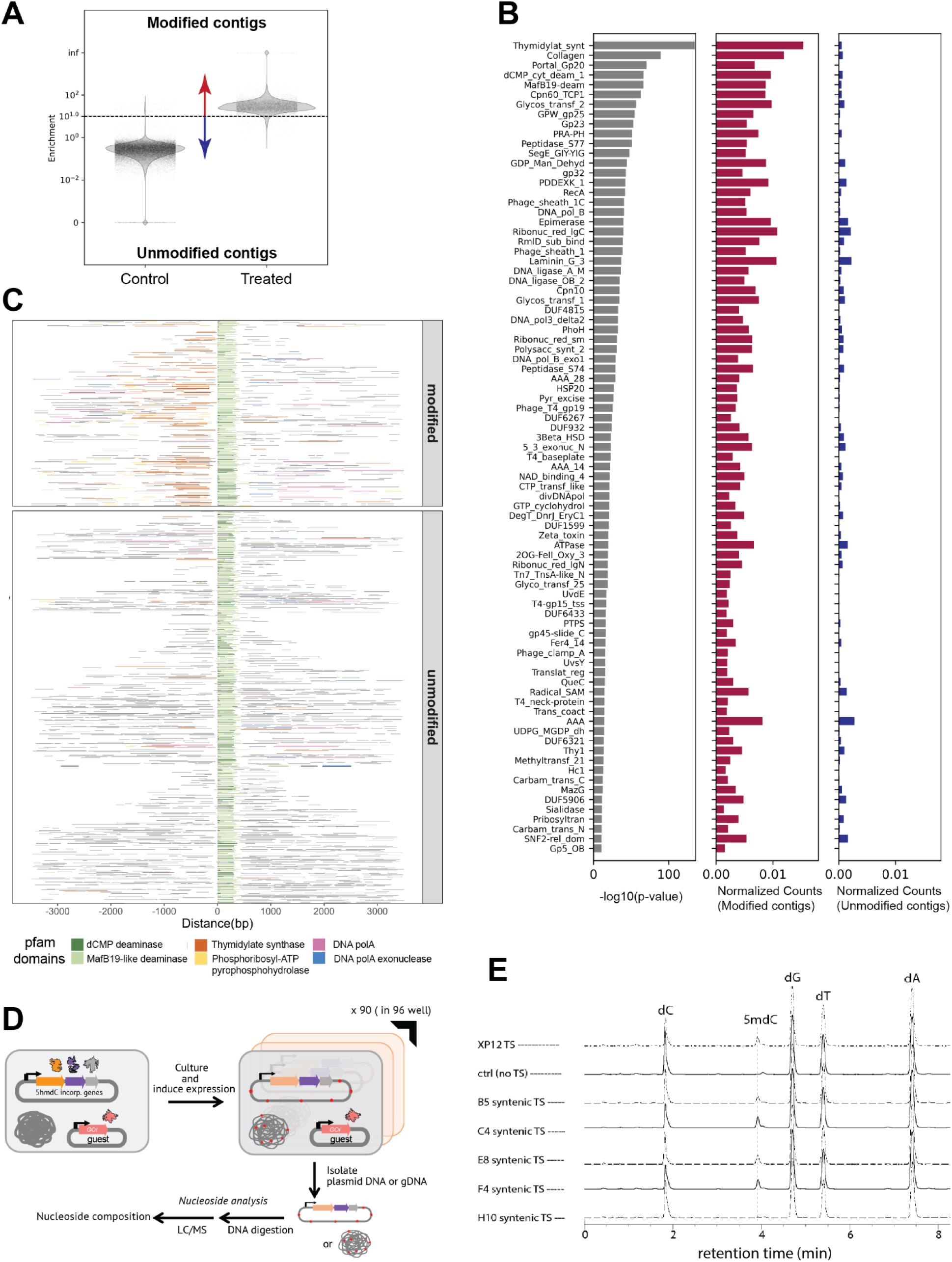
Identification of dCMP deaminases in metaviriome. **A)** Enrichment scores for contigs from the control (left) and treated (right) samples. An enrichment score of 10 or above was used to predict contigs with cytosine modification. Contigs with unmodified cytosine have an enrichment score < 10. **B)** Significantly enriched Pfam annotations (log10(p-value) <= −10) identified in MetaGPA workflow ranked by ascending p-values (Left panel) with their normalized count in contigs with predicted modified (center panel) and canonical C (right panel). **C**) Genes flanking the dCMP deaminases within a maximum of +/− 3kb genomic region. All annotated dCMP deaminases identified from MetaGPA datasets are included in this plot. Color bars represent the top five domains co-localized with dCMP deaminase, namely thymidylate synthase (thymidylat_synt), MafB19 like deaminase (MafB19-deam), Phosphoribosyl-ATP pyrophosphohydrolase (PRA-PH), DNA polymerase A (DNA_pol_A) and 3’-5’ exonuclease (DNA_pol_A_exo1). **D)** Workflow of *in vivo* expression and screening strategy to validate methylation selective deaminase. **E**) LC-MS traces showing biosynthetic activity of deaminase-associated thymidylate synthases (TS). Production of 5mdC has been confirmed for five of the *thyA* homologs recovered from MetaGPA screens, as well as the by the thymidylate synthase homolog from *Xanthomonas* phage Xp12.

**Supplementary Figure 2:**
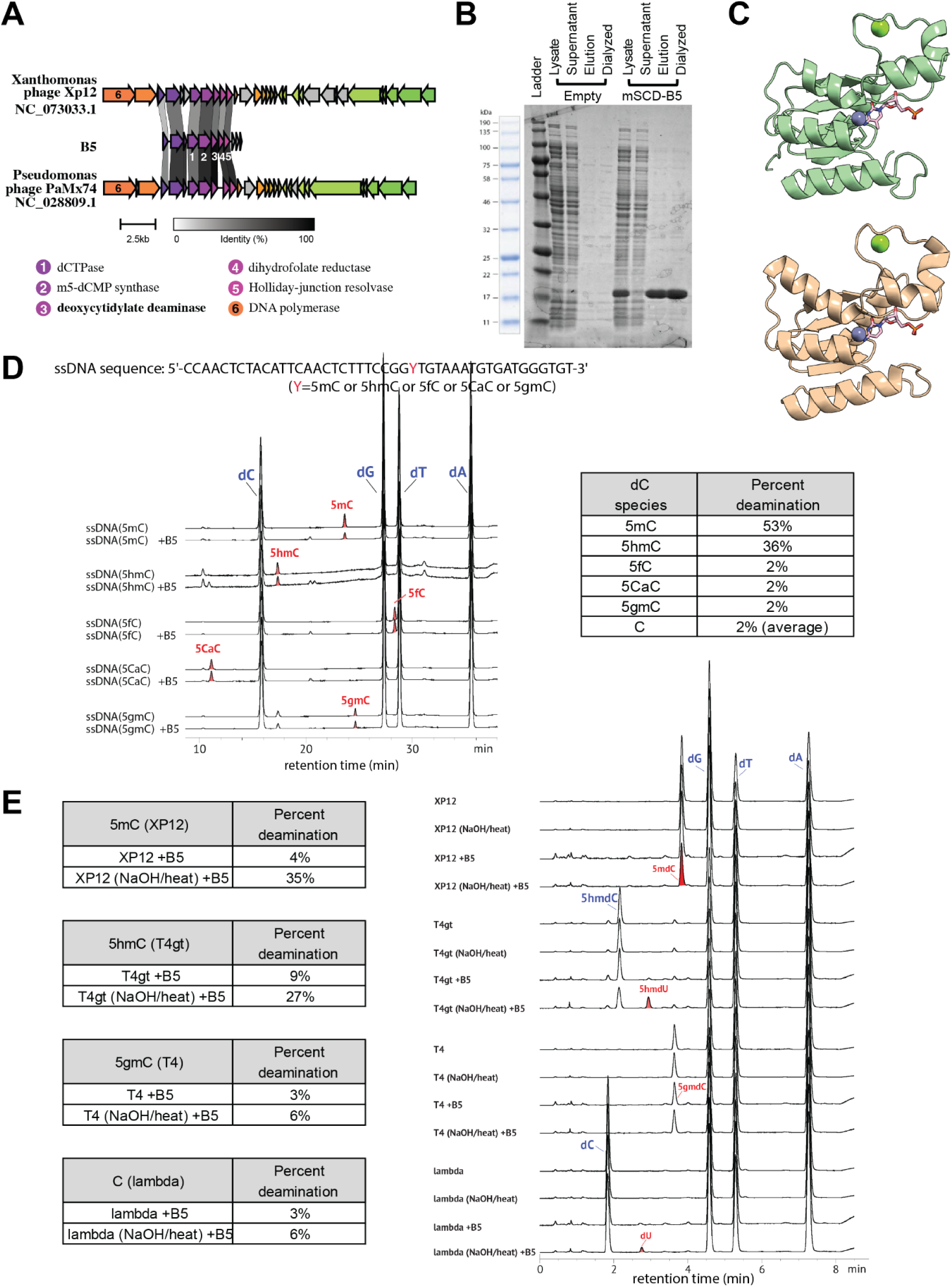
Characterization of the mSCD-B5 deaminase and its deamination rate *in vitro*. **A)** Clinker (Gilchrist and Chooi 2021) schematic showing the deaminase genomic neighborhood for phage Xp12, the contig encoding mSCD-B5, and phage PaMx74. **B**) SDS-PAGE gel showing mSCD-B5 expression and purification. An empty plasmid experiment was included as control. **C**) Alphafold (Tunyasuvunakool et al. 2021) rendering of the mSCD-B5 representative dCMP deaminase (green, upper panel) and its closest Foldseek (van Kempen et al. 2024) structural match in the PDB (wheat, lower panel; 7FH4, Chlorovirus PBCV-1 bi-functional dCMP/dCTP deaminase). **D**) deamination assay monitored by LC-MS on oligonucleotides containing internal cytosine modifications. Quantification of deamination is shown in the right panel ss: single strand, ds: double strand, mC: 5-methylC, hmC: 5-hydroxymethylC, fC: 5-formylC, caC: 5-carboxyC, gmC: 5-glucosyl-methylC. **E**) LC-MS traces (left) and result table (right) of deamination on double strand (-NaOH) or denatured (+NaOH) double strand genomic DNA.

**Supplementary Figure 3:**
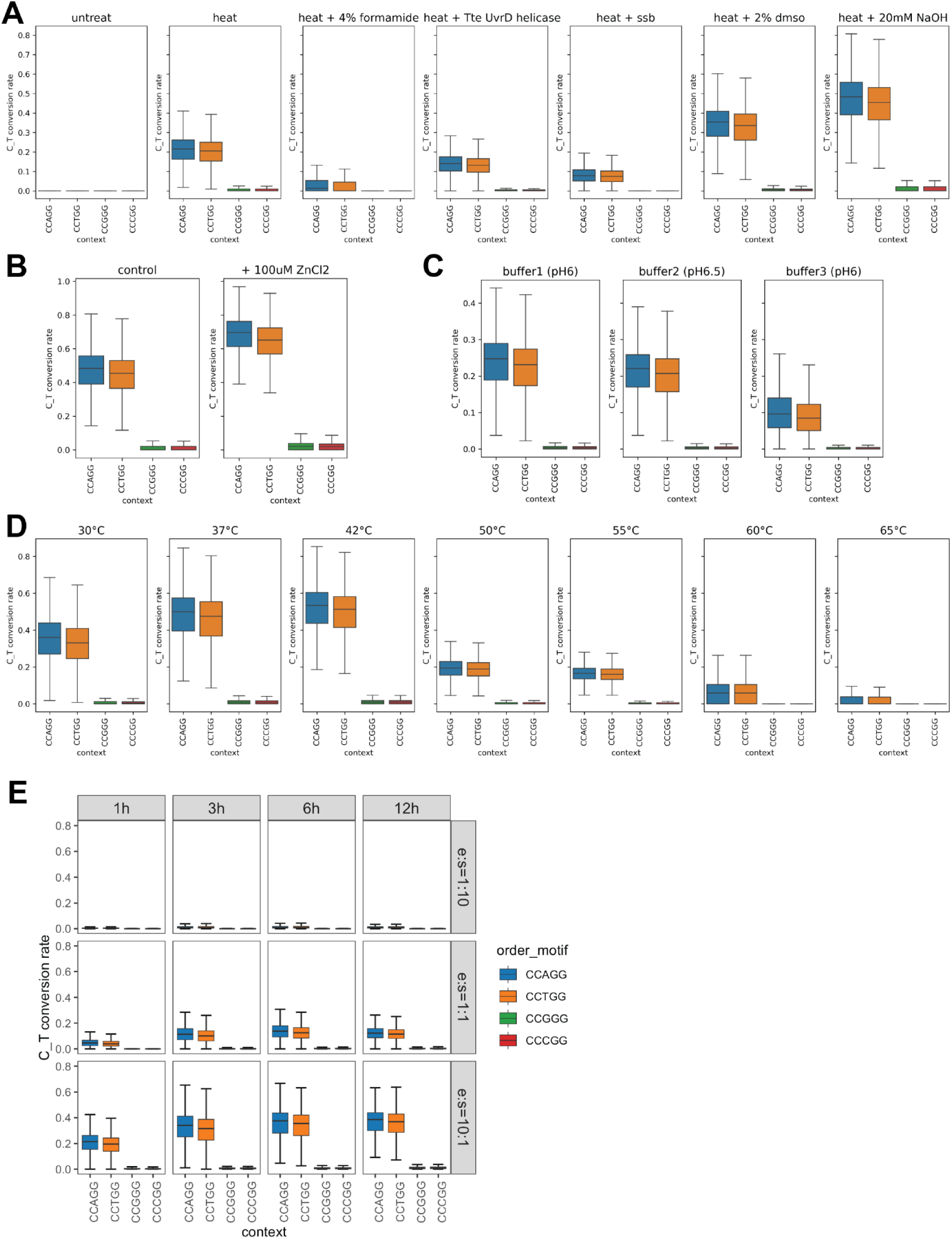
Optimization of high throughput sequencing of mSCD-B5 treated libraries. **A**) The influence of buffer composition during the denaturation step was evaluated by analyzing the fraction of C-to-T substitution (the deamination rates with 1 correspond to 100%) at every C<u>C</u>NGG site in the E. coli DHB4 genome. Specifically, deamination rates were assessed for unmethylated cytosines within C<u>C</u>SGG contexts (where S = C or G) and for methylated cytosines within C5m<u>C</u>WGG contexts (where W = A or T). Conditions tested included heat alone and heat combined with 4% formamide, *Tte* UvrD helicase (NEB #M1202), single-stranded DNA binding protein, 2% DMSO, or 20 mM NaOH. Ssb: single strand DNA binding protein, T4 gene 32 protein (NEB #0300) was used in this study. **B)** Effect of Zn^2+^ in reaction. 100µM ZnCl_2_ was added to the deamination reaction. **C)** Effect of pH buffer in reaction. Buffer 1: Deamination Reaction Buffer (NEB, #E3356); buffer 2: Bis-Tris pH6.5; buffer 3: Bis-Tris pH6 + 100mM NaCl + 10mM MgCl_2_. **D)** Effect of incubation temperature from 30 degree C to 65 degree C. **E)** Effect of enzyme:substrate (E:S) ratio (from E:S=1:10, top to E:S=10:1, bottom) and incubation time (from 1 hour, left to 12 hours, right).

**Supplementary Figure 4:**
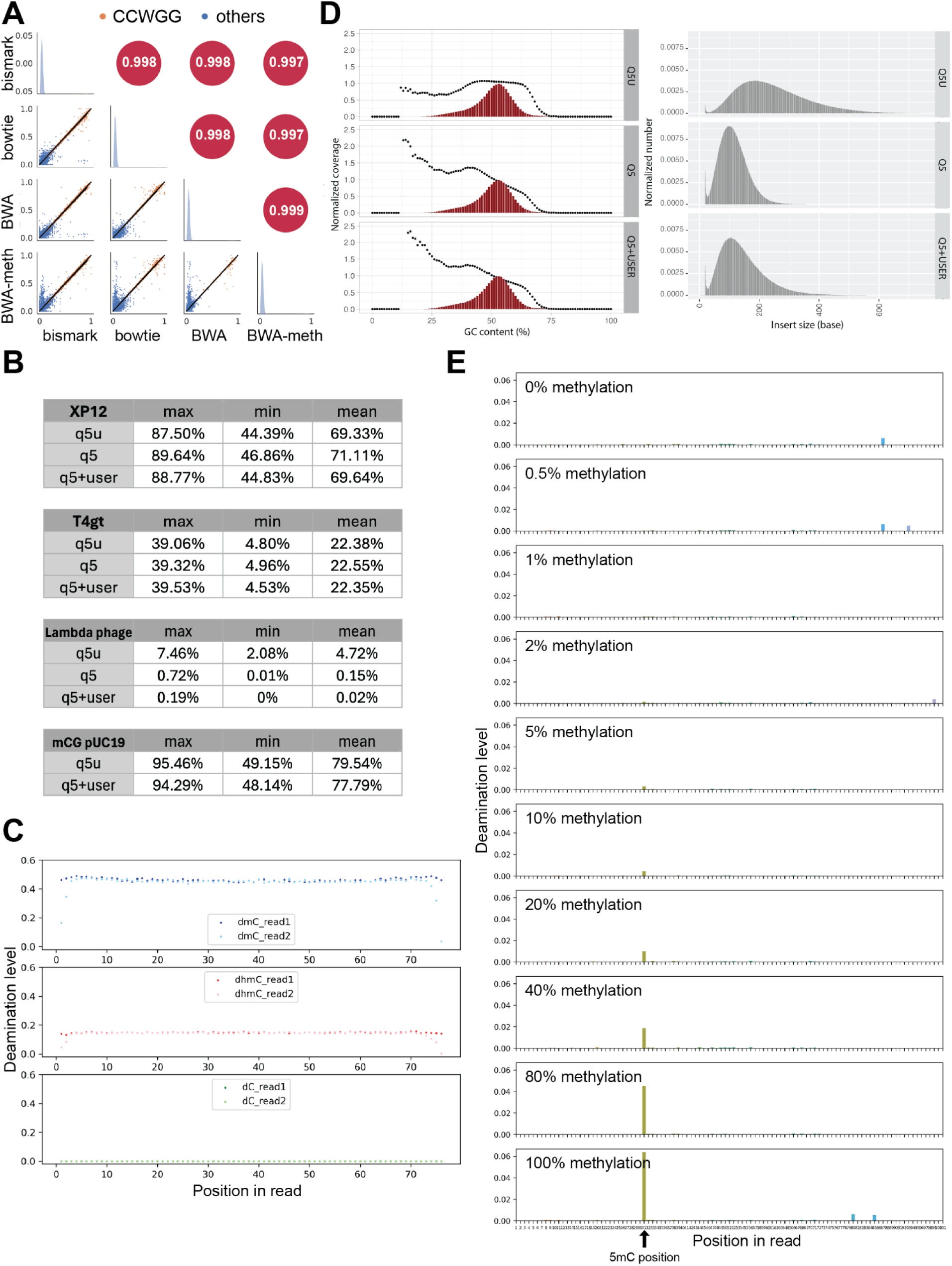
**A**) Comparison of four commonly used mapping tools in methylation detection sequencing analysis. Regression plots in the lower triangle show correlations of deamination level in dcm+ *E. coli* DHB4 per base resolution between tools. Upper triangles show Pearson’s coefficients of detected deamination level between each pair. Distributions of data are shown in diagonal. **B**) Table shows maximum, minimum and mean deamination levels of 5mC (XP12), 5hmC (T4gt) and C (lambda) in NNCNN contexts**. C)** Deamination rate relative to position on the read for 5mC (top), 5hmC (middle) and canonical C (bottom) **D)** Normalized coverage on the E.coli genome function of the GC content (in %, data obtained using Picard tools CollectInsertSizeMetrics). The red bar plots represent the GC distribution expected across the E.coli genome. Size distribution (in bases) of the sequenced libraries amplified by Q5U (top), Q5 (center) and Q5+USER (bottom). Data obtained using Picard tools CollectGcBiasMetrics. **E**) Deamination level relative to position in synthetic oligo carrying different methylation levels. The internal 5mC position is marked with a black arrow.

**Supplementary Figure 5:**
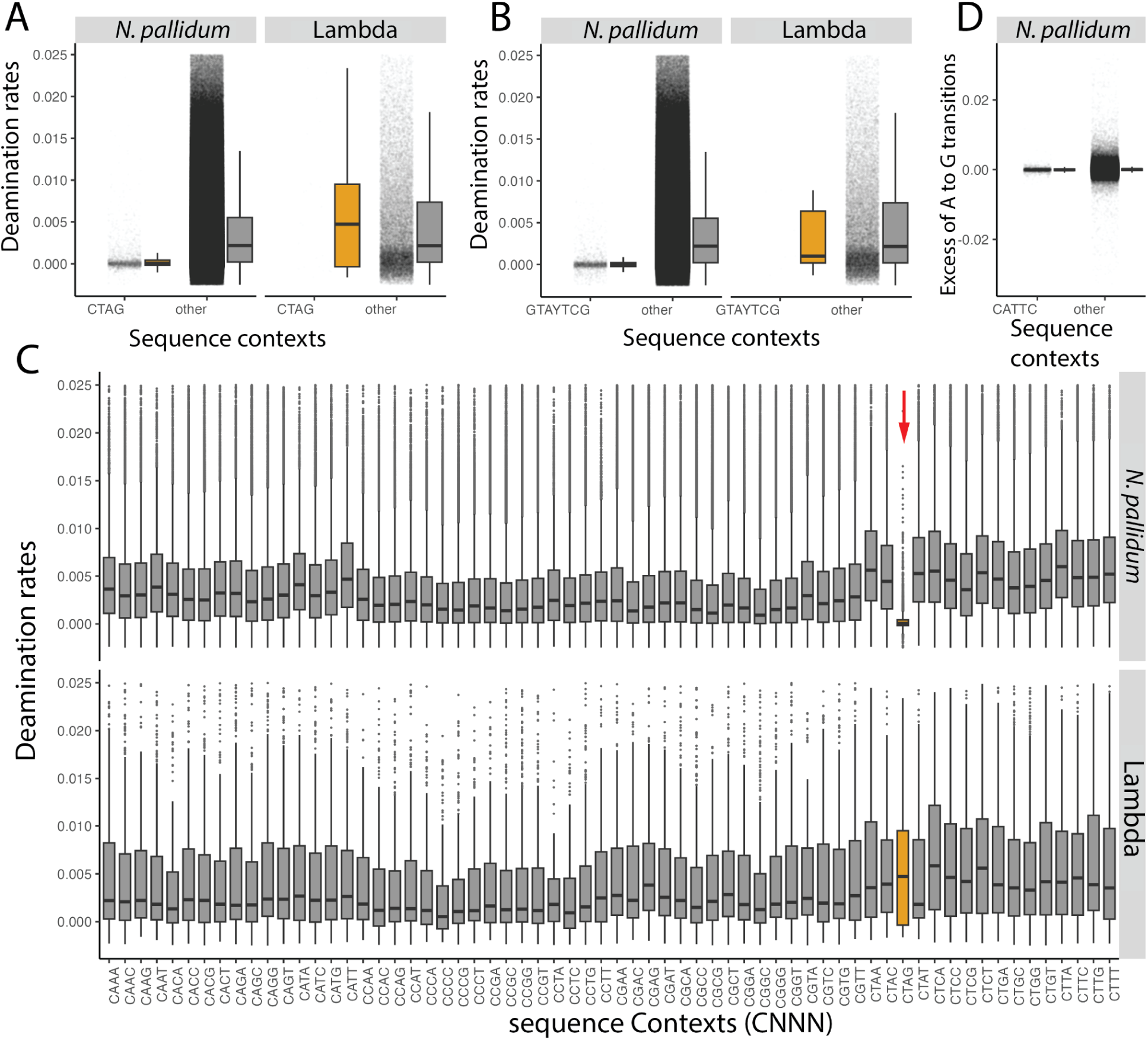
m4C and m6A are not substrates of mSCD represented by mSCD-B5. **A**) Dotplots and boxplots of deamination rates at <u>C</u>TAG sites (left plots) and other sequence contexts (right plots) for *N. pallidum* (left panel) and Lambda (right panel) genomes. **B)** Same as **A)** for the GTAYT<u>C</u>G sites. **C)** boxplots of deamination rates at every CNNN sites in *N. pallidum* (top panel) and Lambda (bottom panel) genomes. **D)** Dotplots and boxplots of excess of A-to-G substitution at C<u>A</u>TTC sites (left plots) and other sequence contexts (right plots) for *N. pallidum.* Excess of A-to-G represents the difference between A-to-G in read1 and read2 of paired-end reads. Average deamination rate for N. pallidum is 0.000142 at GTAYT<u>C</u>G sites (with Y=C or T), 0.000384 at CTAG sites and 0.0383 for all other contexts. In contrast, the average deamination rate for Lambda is 0.00311 at GTAYT<u>C</u>G sites and 0.0117 at <u>C</u>TAG sites.

**Supplementary Figure 6:**
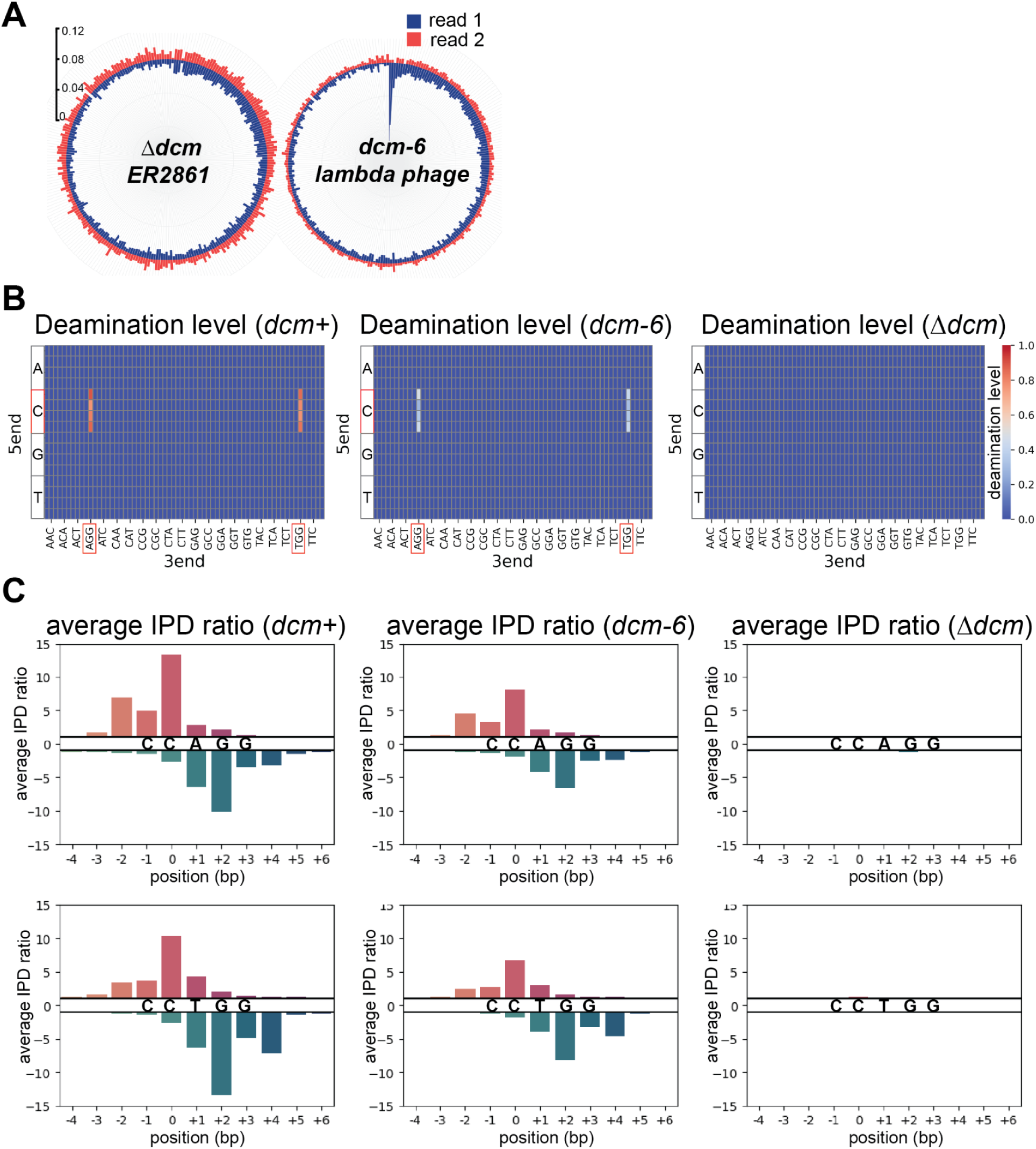
Detection of residual methylation at CCWGG or *dcm*+, *dcm-6* and *Δdcm E.coli* strain. **A)** Circle plot of the average C-to-T transition rate (in %) for all NCNNN sequence contexts in the first of paired reads (R1, blue) and second of paired reads (R2, red) in E. coli ER2861 (Δdcm, left) and Lambda phage grown in E.coli (dcm-6, right). Sequence contexts were ordered by transition rate in read 1 descending order. The 2 contexts with the highest transition rate in R1 were C<u>C</u>TGG and C<u>C</u>AGG (corresponding to C<u>C</u>WGG, dcm site). **B**) Heatmap of deamination levels in NNCNNN contexts (with N=A,T,C or G) in *dcm+* strain (left), *dcm-6* strain (middle) and *Δdcm* strain (right). X axis represents the 2-bases contexts 3’ to the interrogated cytosine, Y axis represent the 3-bases contexts 5’ to the interrogated cytosine and the z-axis (color) represents the average deamination level. Color scale ranges from 0.0 to 1.0 (100% deamination). Deaminated contexts are highlighted as red boxes. **C**) Tet-assisted PacBio sequencing showing average IPD ratio at C<u>C</u>AGG (top) and C<u>C</u>TGG (bottom) for *dcm+* (left), *dcm-6* (middle) and *Δdcm* (right) strains. Position (in base) is relative to the interrogated cytosine at the second base on the C<u>C</u>WGG motif. Positive (red) and negative (blue) values represent average iPD ratios in + and - strand respectively.

**Supplementary Figure 7:**
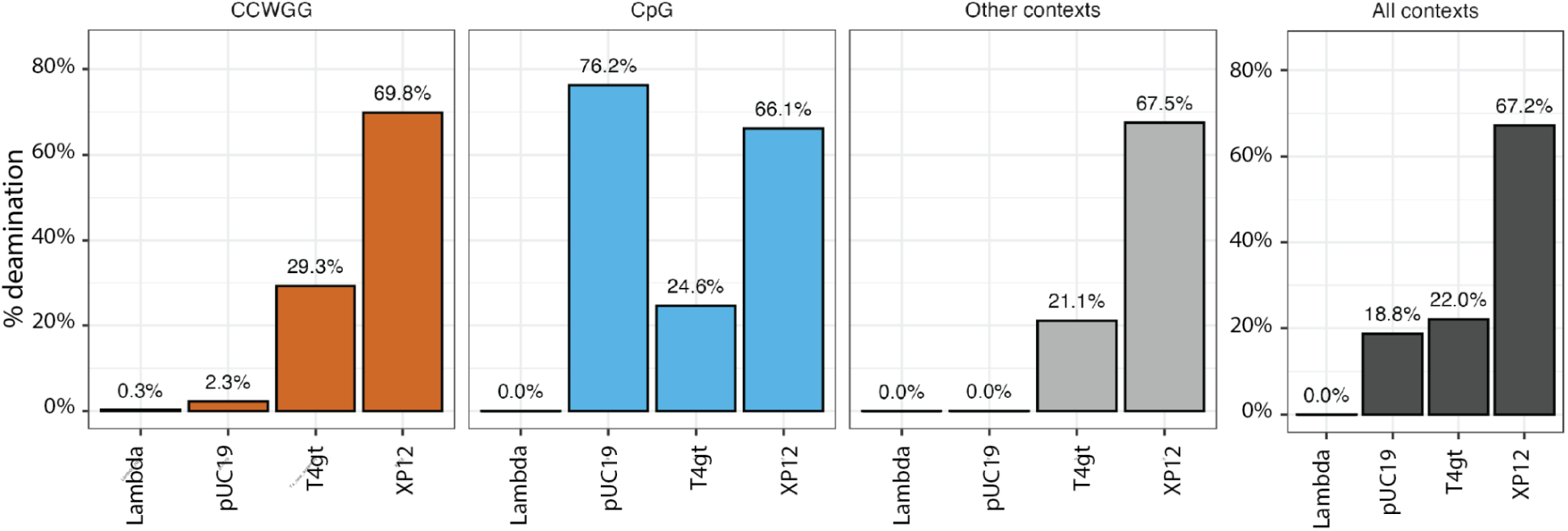
Deamination levels measured from mSCD-B5 treated libraries in control genomes (Lambda, T4gt and XP12) and pUC19 plasmid (5mCpG) in CCWGG contexts (left), CpG context (center left), other NCNNN contexts (center right) and all NCNNN context combined (right).

**Supplementary Figure 8:**
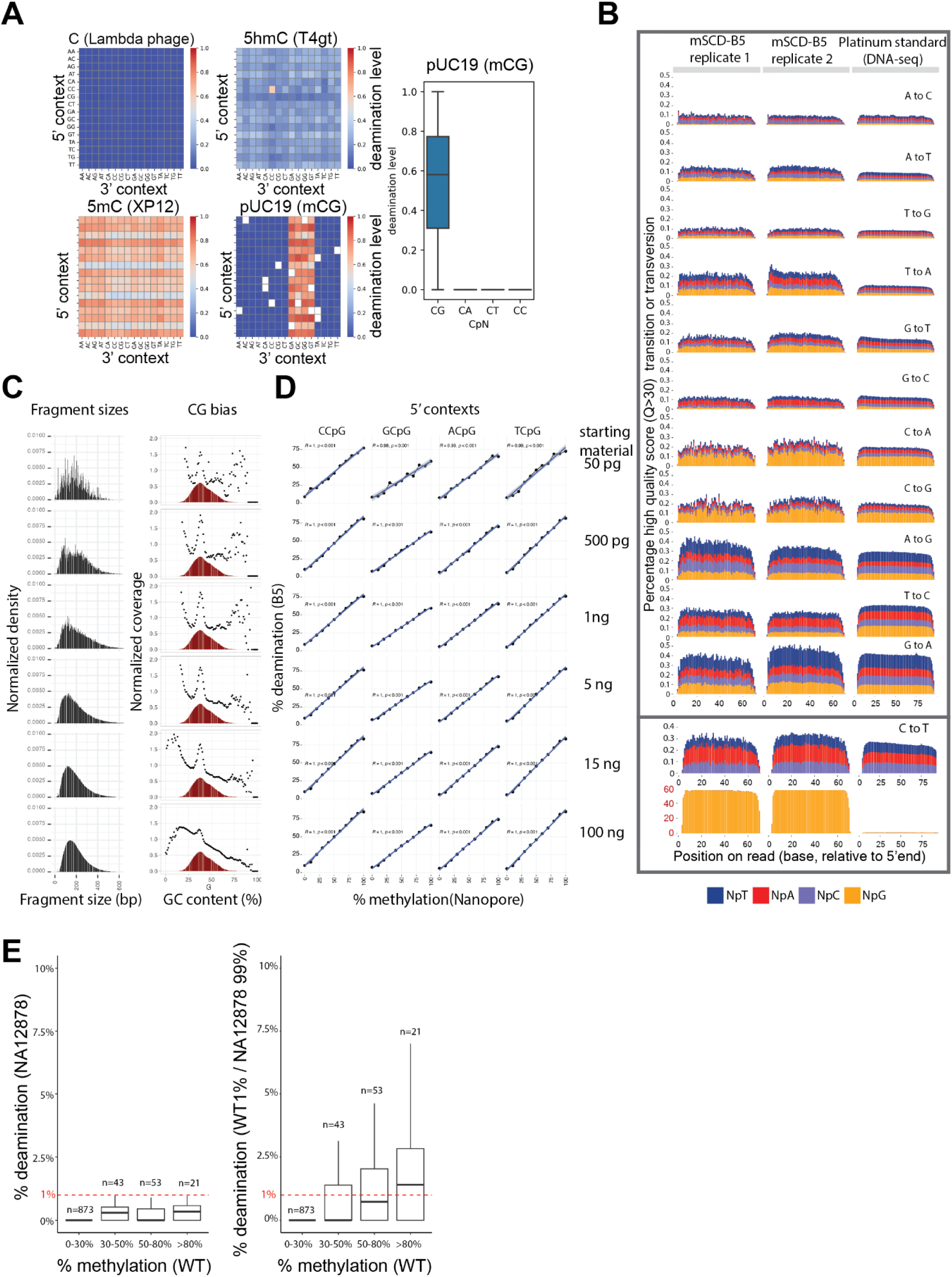
Sequencing of NA12878 genomic DNA treated with mSCD-B5. **A)** Deamination heatmap in spike-in phage genomes or CG-methylated plasmid pUC19. Unavailable sequence contexts in the pUC19 genome were marked blank. Color scale ranges from 0.0 to 1.0 (100% deamination). Deamination level in CN context in CG-methylated pUC19 plasmid DNA is shown in the right panel. The average deamination percentage in CpG context was 76.1%. **B)** Percentage of base substitutions in sequencing reads across read positions (x-axis, in base pairs) is shown for two replicates of mSCD-B5-treated libraries (NA12878 genomic DNA, 100 ng input; left two panels) compared to the platinum standard DNA-seq (NA12878). Substitutions were categorized into 12 types, with a specific panel highlighting C-to-T substitutions in CpG contexts to illustrate the magnitude difference compared to other contexts. This breakdown underscores the distinct patterns of C-to-T conversions in CpG (overwhelmingly due to deamination) versus non-CpG regions (from sequencing error or true variants). Only paired-end read 1 were used to avoid the reverse complement of C-to-T (G-to-A) to interfere with baseline substitution **C)** QC metrics of the fragment size distribution (left panel), GC bias (middle panel) and correlation of the overall-genomic methylation levels obtained with mSCD-B5 (y-axis) and Nanopore (x-axis) at various NCpG contexts (Right panels) for various amount of starting material. **D**) Genome-wide methylation level correlation between Nanopore and mSCD-B5 from 0.05 ng to 100 ng starting material. Nanopore base resolution methylation levels were binned into 0-10%, 10-20% until 90-100% (x-axis) and the equivalent methylation from mSCD-B5 were computed and plotted (Y axis) according to the NCpG context (with N=A,T,C or G). **E)** Methylation levels in CpG island for NA12878 (Left panel) and mixed population of genomic DNA from NA12878 and WT-DKO at ratio 99:1, (Right panel) genomic DNA function of the methylation levels in WT-DKO.

**Supplementary Figure 9:**
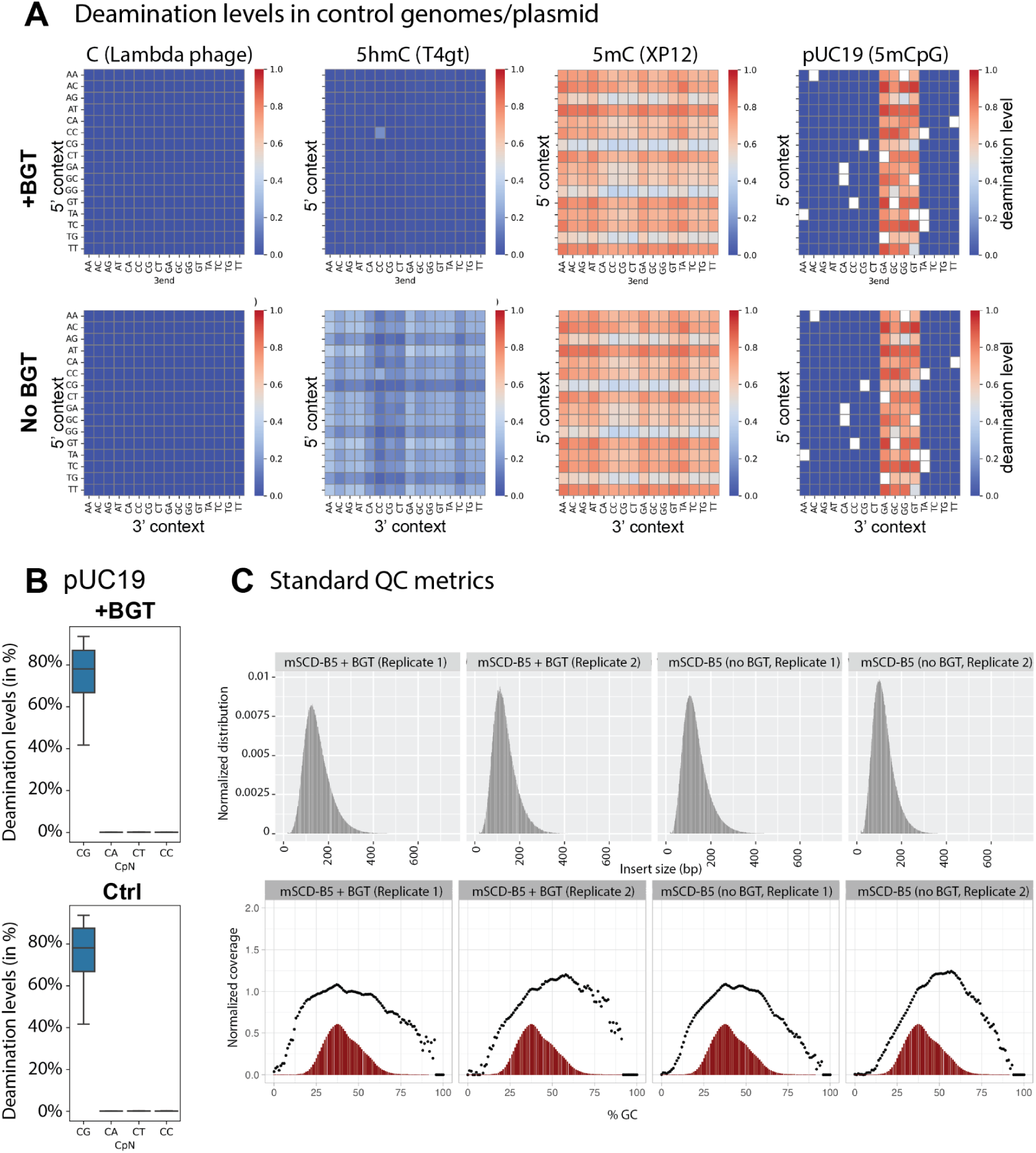
mSCD-B5 deamination analysis in human brain genomic DNA. **A)** Deamination heatmap in spike-in phage genomes or CG-methylated plasmid pUC19. Unavailable sequence contexts in the pUC19 genome were marked blank. Color scale ranges from 0.0 to 1.0 (100% deamination) **B**) Deamination level in all CN contexts for 5mCpG-methylated pUC19 for BGT treated library (+BGT, Top) and no BGT treatment (-BGT, bottom). The average percentage deamination in CpG context in BGT treated and control samples were 74.7% and 74.8% respectively. **C**) Quality control metrics of mSCD-B5-treated libraries derived from human brain genomic DNA. Insert size distribution (in bp, top panels) and GC content bias (bottom panels) for mSCD-B5 + BGT treated libraries (duplicates, left panels) and mSCD-B5 treated libraries (duplicates, right panels). Insert size and GC bias metrics were generated using Picard tools (version 2.0.1)

**Supplementary Figure 10:**
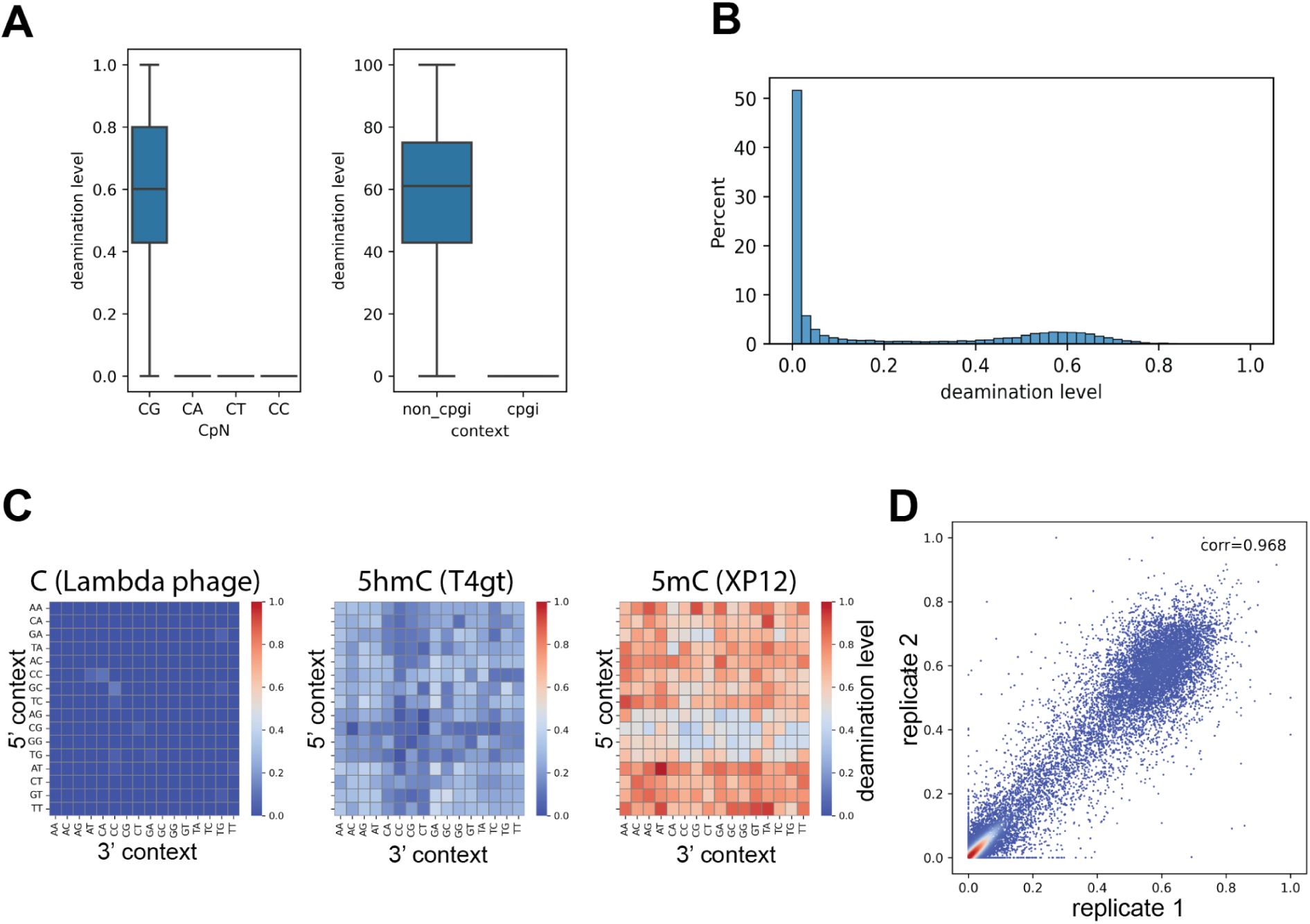
mSCD-seq performed on circulating cell-free DNA. **A)** Detection of methylation on CpN context (left panel) and CpG island context (right panel). **B**) Bimodal distribution of methylation on CpG islands. **C**) Phage spike-in controls demonstrated consistency of selective deamination on mC/hmC. **D**) Correlation plot of deamination levels on CpG islands between two technical repeats. Pearson’s coefficient = 0.968.

